# Essential Gene Phenotypes Reveal Antibiotic Mechanisms and Synergies in *Acinetobacter baumannii*

**DOI:** 10.1101/2022.11.09.515830

**Authors:** Ryan D. Ward, Jennifer S. Tran, Amy B. Banta, Emily E. Bacon, Warren E. Rose, Jason M. Peters

## Abstract

The emergence of multidrug-resistant Gram-negative bacteria underscores the need to define genetic vulnerabilities in relevant pathogens. The Gram-negative pathogen, *Acinetobacter baumannii*, poses an urgent threat by evading antibiotic treatment through both intrinsic and acquired mechanisms. Antibiotics kill bacteria by targeting essential gene products, but antibiotic-essential gene interactions have not been studied systematically in *A. baumannii*. Here, we use CRISPR interference (CRISPRi) to comprehensively phenotype *A. baumannii* essential genes. We show that certain essential genes are acutely sensitive to knockdown, providing a set of promising therapeutic targets. Screening our CRISPRi library against last-resort antibiotics revealed essential pathways that modulate beta-lactam resistance, an unexpected link between NADH dehydrogenase function and polymyxin killing, and the genetic basis for synergy between polymyxins and rifamycins. Our results demonstrate the power of systematic genetic approaches to identify weaknesses in Gram-negative pathogens and uncover antibiotic mechanisms that better inform combination therapies.

**Significance:** *Acinetobacter baumannii* is a hospital-acquired pathogen that is resistant to common antibiotic treatments. *A. baumannii* infections, we need to identify promising therapeutic targets and effective antibiotic combinations. Here, we characterize genes critical for *A. baumannii* viability and their interactions with antibiotics. We find that genes involved in proton gradient formation required for oxygen-dependent energy generation are central to *A. baumannii* physiology and represent appealing drug targets. We show that polymyxins interact with proton gradient genes, explaining how these antibiotics inhibit growth at sub-lethal concentrations and their efficacy in combination therapies. Our studies reveal antibiotic-gene interactions in *A. baumannii* that can inform future therapies.

## Introduction

Gram-negative pathogens, including *Acinetobacter baumannii*, have become increasingly recalcitrant to antibiotic treatment. The emergence of antibiotic resistance alongside a stalled pipeline for additional classes of Gram-negative targeting drugs threaten to precipitate a global health crisis in which many infections become untreatable (1). *A. baumannii* causes serious infections in hospitalized patients and is considered an urgent threat for its ability to evade targeting by antibiotics of last resort (2). *A. baumannii* has numerous defenses against antibiotics including a propensity to acquire resistance genes through horizontal transfer (3, 4), low membrane permeability coupled with robust efflux to prevent antibiotics from reaching cytoplasmic targets (5), and rapid accumulation of resistance mutations (6). Although the unique strengths of *A. baumannii* in resisting antibiotic killing have been well documented, less is known about whether *A. baumannii* carries any unique weaknesses that could be therapeutically exploited.

Comparing *A. baumannii* to model Gram-negatives, such as *Escherichia coli*, has limited utility due to important differences in physiology. In contrast to other Gram-negative ESKAPE pathogens (*Klebsiella, Pseudomonas*, and *Enterobacter*), *A. baumannii* is an obligate aerobe that requires oxidative phosphorylation to generate ATP (7). The outer membrane of *A. baumannii* contains lipooligosaccharide (LOS), rather than lipopolysaccharide (LPS); LOS contains lipid A and core oligosaccharide moieties, but lacks repeating units of O-polysaccharide (8). Critically, LOS is not essential in certain *A. baumannii* strains, including clinical isolates; LOS^-^ strains cannot be targeted by polymyxin antibiotics that bind to lipid A (9, 10). Finally, *A. baumannii* has numerous genes of unknown function, including essential genes that are not present in model bacteria (11, 12). These distinctions underscore the importance of examining antibiotic-gene interactions directly in *A. baumannii*.

Systematic studies of *Acinetobacter* species have provided valuable physiological insights, although *A. baumannii* essential genes have not been comprehensively characterized. Transposon sequencing (Tn-seq) studies in *A. baumannii* have identified putative essential genes (11, 12), cataloged antibiotic-gene interactions for non-essential genes (13, 14), defined the functions of uncharacterized genes (14), and uncovered the mechanism for strain-specific essentiality of LOS biosynthesis (15). An elegant Tn-seq study of non-pathogenic *Acinetobacter baylyi* monitored depletion of strains with disrupted essential genes following natural transformation (16), but it is unclear whether those findings are directly applicable to *A. baumannii*. CRISPR interference (CRISPRi (17)) can be used to phenotype essential genes via partial knockdown or depletion time course, but CRISPRi has only been applied to a handful of *A. baumannii* essential genes so far (12).

To systematically probe for weaknesses in *A. baumannii*, we generated and screened a pooled CRISPRi library targeting all putative essential genes (Fig. 1A). We identified essential genes that are most vulnerable to depletion, thereby prioritizing targets for future drug screens. We further used CRISPRi to define the mode of action for antibiotics of last resort, finding antibiotic targets and genetic barriers to drug efficacy, as well as the genetic bases for synergistic antibiotic combinations.

**Fig. 1.**
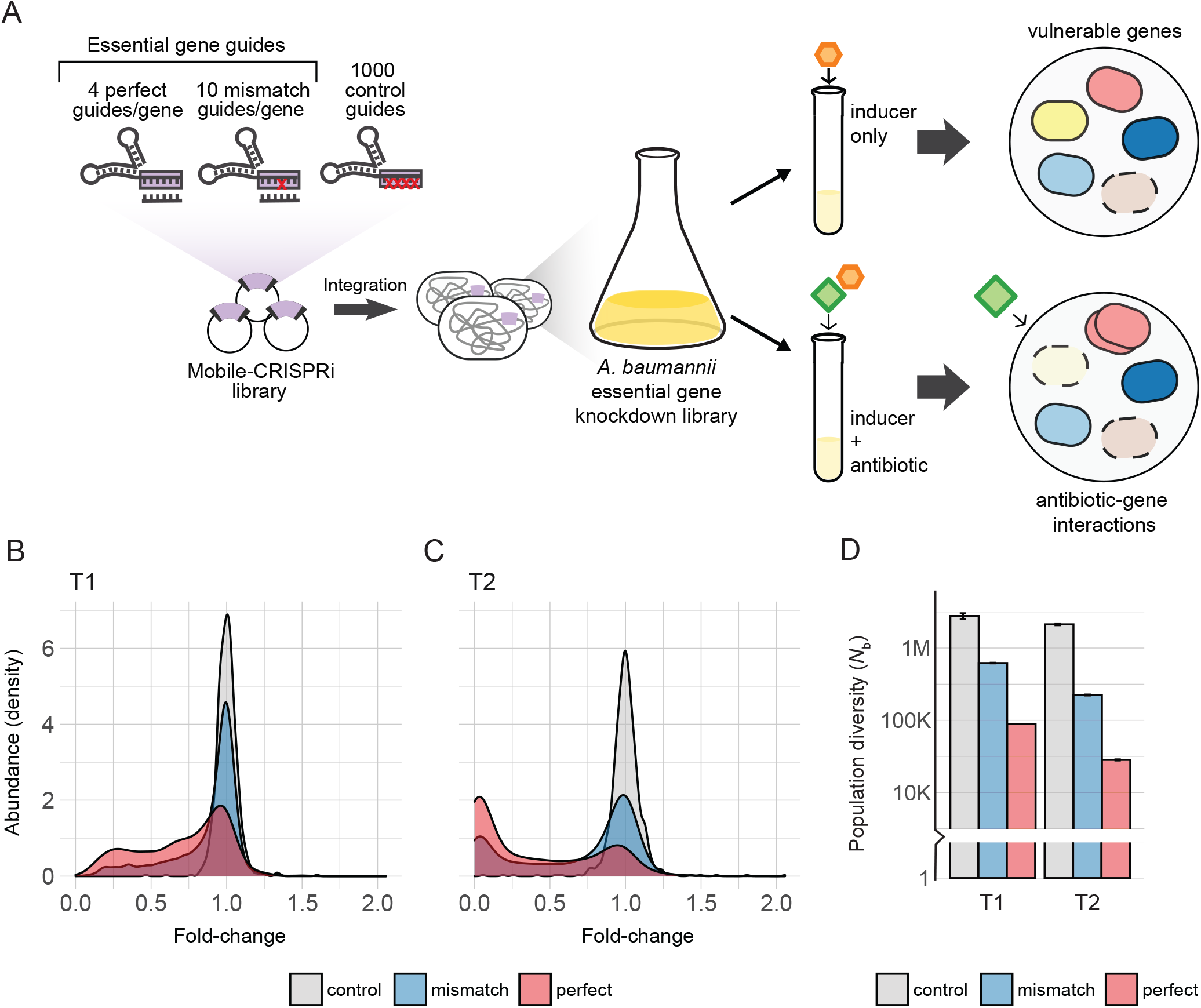
*A. baumannii* essential gene CRISPRi library. A) 4 perfect and 10 mismatch guides per essential gene and 1000 non-targeting (control) guides were cloned into Mobile-CRISPRi plasmids with inducible dCas9 and sgRNA and transferred to *A. baumannii*. The pooled library was then treated with inducer to assess gene vulnerability or with inducer and antibiotic to examine antibiotic-gene interactions. B) Relative abundance or kernel density estimation (compared to T0) of control, mismatch, and perfect guides across fold-change at T1 and C)T2. D) Bottleneck number (*N*_b_) comparing population diversity of control, mismatch, and perfect guides from T0 to T1(left) or to T2 (right).

## Results

### Construction and validation of an *A. baumannii* essential gene CRISPRi library

We first optimized CRISPRi in *A. baumannii*, finding that reduced expression of *dcas9* lowered toxicity and still achieved ∼20-fold knockdown (Supplemental Methods, Fig. S1A-E, and Tables S1-4). We next designed and constructed a CRISPRi library targeting all putative essential genes in the *A. baumannii* type strain 19606 (18). We combined a published Tn-seq analysis of *A. baumannii* AB5075 (11) with ortholog analysis to define a list of CRISPRi target genes we call the “*Ab* essentials” (406 genes total, Table S5, S8). We then designed a computationally optimized CRISPRi library targeting the *Ab* essentials that consisted of three types of single guide RNAs (sgRNAs): 1) perfect match sgRNAs (19) to maximize knockdown (∼4/gene), 2) mismatch sgRNAs (20) to generate a gradient of partial gene knockdowns (∼10/gene), and 3) control sgRNAs that are non-targeting (1000 total). The library was cloned and site-specifically integrated into the 19606 genome using Mobile-CRISPRi (21). Illumina sequencing of integrated sgRNA spacers confirmed that our CRISPRi library targeted all the *Ab* essentials (median = 14 guides/gene; Fig. S2).

Finally, we validated our *A. baumannii* CRISPRi library by measuring depletion of essential gene targeting sgRNAs during pooled growth. We grew the library to exponential phase in rich medium (LB) without induction (T0), diluted back into fresh medium with saturating IPTG to induce CRISPRi and grew cells for ∼7 doublings (T1), then diluted back a second time in IPTG-containing medium and grew cells for an additional ∼7 doublings (T2). To quantify depletion, we calculated fold change (FC) and strain diversity (*Nb*; (22)) of sgRNA spacer counts between T0 and T1 or T2 (Fig. 1B-D, Table S6). Both FC and *Nb* showed depletion of essential gene targeting sgRNAs by T1 and additional depletion by T2, whereas control sgRNAs were largely unaffected at both time points. The lack of an induction effect on control strain abundance suggests that toxic guide RNAs such as “bad seeds” (23) are mostly absent from our library. Taken together, our CRISPRi library effectively and comprehensively perturbs essential gene functions in *A. baumannii*.

### Identification of *A. baumannii* essential genes that are sensitive to depletion

Essential gene products that have a strong, negative impact on fitness when depleted, i.e., “vulnerable,” are high-value targets for antibiotic development. CRISPRi allows us to define vulnerable genes by controlling the duration and extent of depletion (20, 24–26). To identify vulnerable genes, we quantified depletion of strains with perfect match guides from the CRISPRi library during growth in rich medium (LB) (Fig. 2A). At T1, 88 genes showed significant depletion (log2FC<-1 and Stouffer’s *p*<0.05), and by T2 an additional 192 genes were depleted (280/406 total or 69%; Table S7). 126 of the genes on our *Ab* essentials list were non-responsive to depletion by T2; these genes either require >20-fold depletion (20), are false positives from the AB5075 Tn-seq analysis (11), or are not essential in 19606. Screening our library in sub-MIC (minimal inhibitory concentration) levels of clinically relevant antibiotics (see below) recovered phenotypes for 74 of the 126 genes that were non-responsive in rich medium, demonstrating that these genes are important for antibiotic mode of action (Fig. S3A). Overall, the vast majority of *Ab* essentials (354/406 or 87%) show significant phenotypes in our CRISPRi screens.

**Fig. 2.**
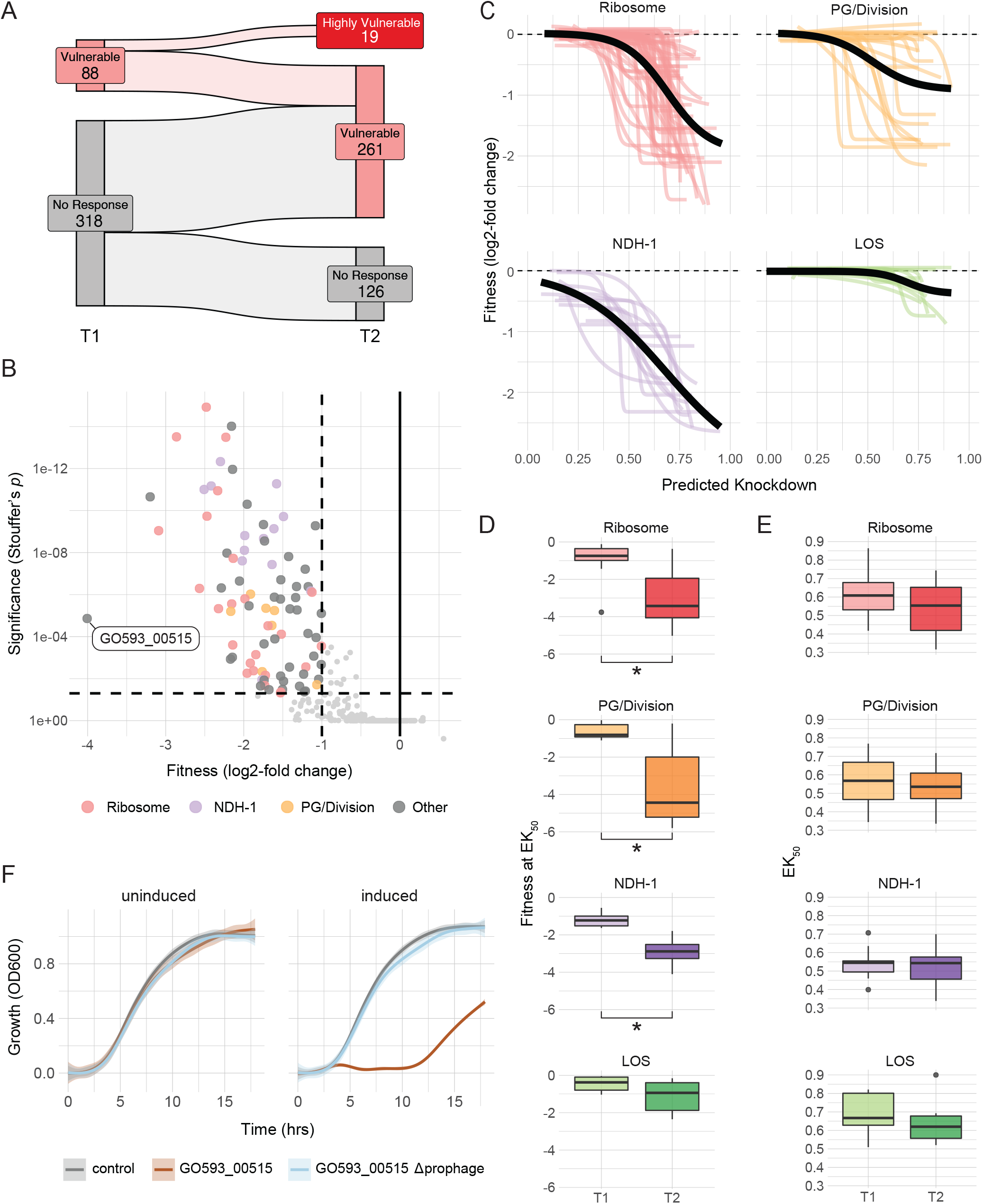
Depletion of *A. baumannii* essential genes. A) Sankey plot of genes at T1 and T2, binned as vulnerable, highly vulnerable, or no response to depletion. Genes are vulnerable if gene-level knockdowns have log2-fold changes < -1 and Stouffer’s p < 0.05. Highly vulnerable genes have log2-fold changes < -8 and Stouffer’s p < 0.05. B) Volcano plot of gene-level knockdowns at T1, plotted by the significance (Stouffer’s *p*) and fitness (median log2-fold change) across perfect guides. Gene-level knockdowns with log2-fold change < -1 and Stouffer’s *p* < 0.05 are highlighted, and those involved in ribosome, NDH-1, and PG/division pathways are in color. No gene-level knockdowns related to LOS met these fitness and significance parameters. C) Knockdown-response curves for ribosome, PG/division, NDH-1, and LOS pathways. Colored curves represent individual genes in each pathway, generated by fitting curves to the fitness (log2-fold changes) of mismatch guides with a range of predicted knockdown for each gene. A predicted knockdown of 1 represents maximal knockdown, equivalent to a perfect guide. Pathway knockdown-response curves, fit to all mismatch guides for all genes in the pathway, are represented with a black line. D) Fitness at EK50 (half-maximal effective knockdown) by pathway. Ribosome, PG/division, and NDH-1 pathways showed a statistically significant (*p* < 0.05) drop in fitness at EK50 between T1 and T2, but LOS did not. Significance was calculated by Mann–Whitney U test, \**p* < 0.001. E) EK50 for ribosome, PG/division, NDH-1, and LOS pathways. There are no significant differences (*p* < 0.05) between T1 and T2 for any pathway. F) Growth, measured by OD600, over 18 hours for a non-targeting control strain, GO593_00515 knockdown strain, and GO593_00515 knockdown strain with a deleted prophage in LB (uninduced, left) or LB with 1 mM IPTG (induced, right).

We first focused on genes that were depleted from the pool at T1 as these genes were likely to be the most vulnerable of the *Ab* essentials (Fig. 2B). As expected, T1 vulnerable genes were enriched for those encoding ribosomal proteins (FDR = 0.0022, KS test), consistent with tight coupling between fitness and ribosome function (27) as well as previous CRISPRi screens targeting essential genes in model bacteria (20, 24, 28–30). In contrast, genes involved in oxidative phosphorylation were uniquely vulnerable in *A. baumannii* (FDR = 2.72e^-06^, KS test); whereas these genes are not essential in the model bacterium, *E. coli* (31). Among the oxidative phosphorylation outliers, genes encoding the NADH dehydrogenase complex I (NDH-1; *nuo* genes) constituted their own enrichment group (FDR = 7.36e^-06^, AFC test). These results highlight the central requirement of proton translocation for ATP generation in an obligate aerobe.

By T2, most *Ab* essentials were depleted (Fig. 2A and S3B), but a set of “highly vulnerable” genes stood out by magnitude of depletion (defined arbitrarily as the top 5%; depleted by >250-fold); these genes were also identified as vulnerable at T1 (Fig. 2A). The highly vulnerable set contained genes involved in peptidoglycan deposition during cell division; six of the top 10 most depleted genes functioned in PG precursor synthesis and deposition (*murG*/*F*/*A, ftsI*) or cell division (*ftsA*/*Z*). Indeed, genes related to cell division were strongly depleted as a group at T2 (FDR = 0.0058, AFC test). Morphological analysis of “terminal phenotypes” caused by essential gene depletions (24) and mechanistic studies of cell-wall-targeting antibiotics (32) have shown that inhibiting *mur* and *fts* gene products results in a loss of cell shape and subsequent lysis, consistent with a sharp drop in abundance for these genes in our pooled library.

To validate and extend our vulnerable gene findings, we next examined depletion across a knockdown gradient using Mismatch-CRISPRi. We previously developed a machine learning approach to predict the knockdown efficacy of mismatched sgRNAs relative to their perfect match parent guides (20). As expected, mismatch guides targeting vulnerable genes with high predicted efficacy showed increased depletion relative to those with low predicted efficacy (Fig. 2C); this pattern was generally consistent across the *Ab* essentials (Fig. S3C), confirming that our machine learning approach can predict mismatch guide behavior in *A. baumannii*. As seen in model bacteria (20) and *Mycobacterium tuberculosis* (33), the relationship between knockdown and fitness varied by pathway and gene (Fig. 2C). To describe the relationship between predicted knockdown and fitness (log2FC), we used an approach developed by Reynolds and colleagues to fit the data with a 4-parameter logistic function (26) producing what we call “knockdown-response curves.” Plotting these curves for individual genes involved in translation (ribosome), cell envelope (PG/division), and oxidative phosphorylation (NDH-1) revealed strong fitness impacts with partial knockdowns, but little impact on non-essential genes such as those involved in LOS biosynthesis. Knockdown-response curves for individual genes in pathways tended to be similar, with some genes in PG/division showing increased vulnerability (e.g., *murG*).

Knockdown-response curves allowed us to determine the extent of knockdown required to generate phenotypes. We defined the level of predicted knockdown required to elicit half of the maximal effect on fitness as the EK50. As expected, the median log2FC at EK50 for vulnerable genes (e.g., those encoding PG/division genes) showed greater depletion at T2 versus T1 (*p* <0.05, t-test (Fig. 2D)). Interestingly, there was no significant difference between median EK50 values at T1 versus T2 for all pathways tested (Fig. 2E); this indicates that the half-maximal knockdown required to elicit a fitness effect generally does not change despite large changes in the maximal fitness effect. We infer that guides causing phenotypes under one experimental condition will tend to accumulate phenotypes under different conditions, while guides with insufficient knockdown to elicit phenotypes will continue to be non-responsive. This finding has implications for designing compact and responsive essential gene knockdown libraries in *A. baumannii* and possibly other organisms. In sum, pathways for oxidative phosphorylation, peptidoglycan biosynthesis, and translation are vulnerable targets in *A. baumannii*.

A gene encoding a putative Arc family transcriptional repressor (GO593_00515) was among the most depleted genes at both T1 and T2. Arc family repressors have been extensively studied for their role in the Phage P22 life cycle (34). As expected, GO593_00515 is located within a predicted prophage in the 19606 genome; this locus lacks a prophage in *A. baumannii* strain 17978 and is occupied by a similar, but distinct prophage in AB5075 (Fig. S4A). Synteny between the 19606 prophage and P22 suggested that GO593_00515 could be involved in lysogeny maintenance. GO593_00515 knockdown cells showed little growth 10 hours after dilution into IPTG-containing medium (Fig. 2F). Addition of IPTG to growing GO593_00515 knockdown cells resulted in lysis after 7 hours (Fig. S4B), although supernatant from lysed cells failed to form detectable plaques on a lawn of 17978. We reasoned that if the essential function of GO593_00515 is to repress expression of toxic prophage genes, we could suppress its essentiality by deleting the prophage entirely. Indeed, IPTG induction of the GO593_00515 knockdown in the presence of an integrated knockout plasmid allowed us to recover prophage deletion strains that also lacked GO593_00515 (Fig. 2F, S4C, and S4D).

Thus, repression of toxic prophage genes is a critical but conditionally essential function in *A. baumannii*. Given the ubiquity of prophages harboring toxic lysis genes (35), we suggest that knockdown of phage repressors could aid in identifying proteins that are exceptional at lysing *A. baumannii*.

### Essential gene knockdowns that potentiate or mitigate carbapenem action in *A. baumannii*

Mode of action studies can determine the direct targets of antibiotics as well as genes in related pathways that positively or negatively impact antibiotic function. To test if our CRISPRi library could define antibiotic function in *A. baumannii*, we screened against imipenem and meropenem,two carbapenem class beta-lactams that are used to treat multidrug-resistant *A. baumannii* strains (36). Beta-lactam antibiotics target penicillin-binding proteins (PBPs), and the essential septal PBP in *A. baumannii* is encoded by *ftsI* (11). We found that *ftsI* was either the most depleted or among the most depleted *Ab* essentials at T1 when cells were grown in sub-MIC doses of imipenem or meropenem (Fig. 3A and S5A). By T2, *ftsI* remained a significant outlier, but depletion of genes in related pathways made it less obvious that *ftsI* was the direct target (Fig. S5B). Strong sensitive outliers at both time points included the *mur* genes, which are involved in synthesizing PG precursors that are cross-linked to the growing sacculus by FtsI during cell division (37). These results demonstrate that CRISPRi can define antibiotic targets and related pathways in *A. baumannii*.

**Fig. 3.**
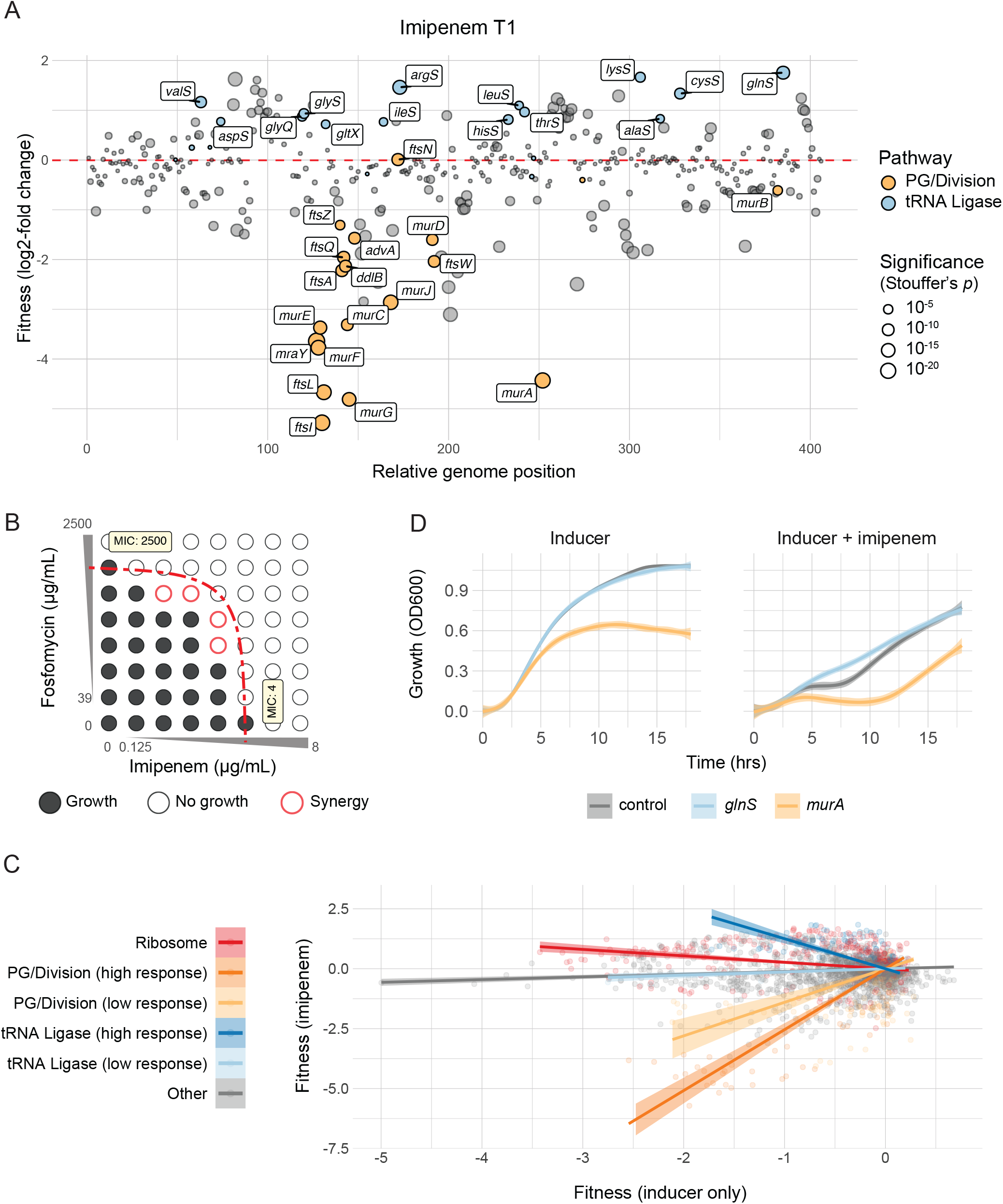
Gene interactions with carbapenems in *A. baumannii*. A) Bubble plots of fitness, or median log2-fold change, for each gene in imipenem (0.09 µg/mL) at T1 compared to the inducer only (no drug) sample. Bubbles are plotted by gene position, or relative location on the genome compared to other predicted essential genes. Significance (Stouffer’s *p*) is represented by size; genes in PG/division (orange) or tRNA ligase (blue) pathways are highlighted. B) Sensitivity assay for fosfomycin-imipenem interactions. 2-fold serial dilutions of drugs from minimum inhibitory concentrations (MICs) represented by gray wedges. No growth below dotted line, fractional inhibitory concentration (FIC) index ≤ 0.5, shows synergy (red border). C) Scatterplot of mismatch guide fitness (log2-fold change) in inducer only compared to relative fitness in imipenem at T1. Lines represent linear model fits with 95% confidence interval. Guides for genes in PG/division or tRNA ligase pathways are divided into groups using hierarchical clustering based on response to imipenem. D) Growth, measured by OD600, over 18 hours for a non-targeting control strain, *glnS* knockdown strain, and *murA* knockdown strain in LB with inducer (1 mM IPTG, left) or LB with inducer and 0.09 µg/mL imipenem (right).

We considered the possibility that sensitive outlier genes other than the direct target may represent targets for antibiotic synergy. The *murA* gene was a strong outlier in both carbapenem screens, and its product is the direct target of fosfomycin. *A. baumannii* is highly resistant to fosfomycin (38); thus, the drug has little clinical use as a monotherapy. To test for an interaction, we measured synergy between imipenem or meropenem and fosfomycin using a checkerboard assay (Fig. 3B and S5C). We found that both carbapenems synergized with fosfomycin, and addition of 0.25 µg/ml imipenem or meropenem was sufficient to lower the MIC of fosfomycin by 4-fold. Although this synergistic concentration of fosfomycin remains too high for clinical use, our experiment serves as a proof of principle that *A. baumannii* CRISPRi screens can predict potential synergies. These results also provide a likely mechanism for the observed synergy between beta-lactams and fosfomycin in fosfomycin-sensitive *Pseudomonas aeruginosa* (39).

We further analyzed our screen data to identify essential pathways that mitigate carbapenem action when perturbed. Previous work hypothesized that beta-lactam resistance and growth are linearly related: as growth slows, resistance increases (40, 41). However, we found little relationship between growth and carbapenem resistance across our CRISPRi library (R^2^ = 0.005, 0.007; slope = 0.13, - 0.05; *p* < 10^−6^), suggesting that slow-growing strains of *A. baumannii* are not necessarily more tolerant to beta-lactam treatment (Fig. 3C and S6A).

Instead, specific genetic pathways showed increased carbapenem resistance. Gene set enrichment analysis identified genes encoding ribosomal proteins as resistant (FDR = 5.44e^-07^, KS test), consistent with antagonism between beta-lactams and ribosome inhibitors observed for other bacteria (42). Genes involved in tRNA charging were among the most resistant outliers (FDR = 2.96e^-06^, KS test; Fig. 3A and 3C); in particular, a highly responsive subset of tRNA ligases showed a negative correlation between growth and beta-lactam resistance (Fig. 3C, S6A, and S6B). We validated our pooled screening results by showing that knockdown of the tRNA ligase, *glnS*, reduces sensitivity to imipenem despite having a negligible effect on growth in the absence of imipenem (Fig. 3D). The non-targeting control strain showed a characteristic “dip” in optical density (OD) after ∼5 hours of growth in imipenem that was absent from the *glnS* knockdown growth curve (Fig. 3D). We speculate that knockdown of tRNA ligase genes induces a cell stress pathway that allows for continuous growth in imipenem; possibly the stringent response (30). We did not observe differential growth of knockdowns and controls at the OD dip in meropenem-treated cells (Fig. S6D), suggesting that additional mechanisms play into resistance. In sum, we found that *A. baumannii* CRISPRi can identify genetic mechanisms of synergy and antagonism.

### The genetic basis for colistin-rifampicin synergy

Colistin and rifampicin synergistically inhibit *A. baumannii* growth by an unknown mechanism (43). We recapitulated this synergy in our screening conditions (LB medium), finding that colistin can increase rifampicin potency by >10-fold (Fig. S7A). To define the mechanisms for colistin-rifampicin synergy, we first screened our CRISPRi library against colistin and rifampicin individually. Colistin, also known as polymyxin E, is an antibiotic of last resort for treating multidrug-resistant *A. baumannii* (10). Colistin binds to the lipid A moiety of LOS and is thought to kill cells by membrane disruption (44). LOS is not essential in many strains of *A. baumannii*, including 19606, and LOS disruption leads to a >500-fold increase in colistin resistance (15). As expected, screening our library against a sub-MIC dose of colistin identified LOS synthesis genes as resistant outliers (FDR = 1.98e^-08^, KS test; (Fig. 4A and 4B). Among the most resistant outliers were *lpxA* and *lpxC*; these genes encode enzymes that catalyze the first two committed steps in LOS synthesis and are found at high frequencies in selections for colistin-resistant mutants (15). LOS genes further downstream in the biosynthetic pathway showed less resistance, possibly due to a tradeoff between surviving colistin and producing toxic LOS intermediates. Surprisingly, knockdown of genes encoding NDH-1 caused heightened sensitivity to colistin (FDR = 1.42e^-06^, AFC test).

**Fig. 4.**
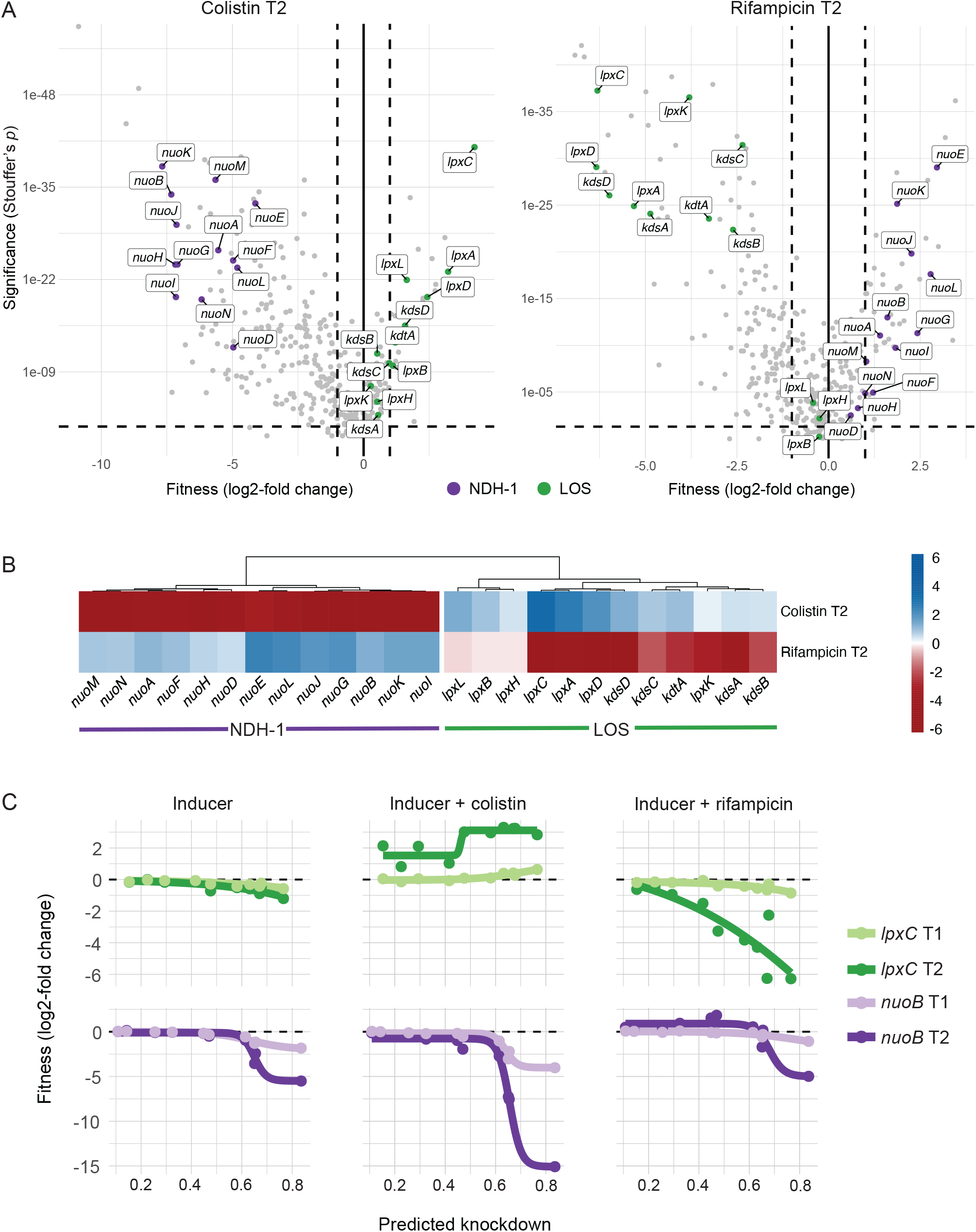
Genetic basis for colistin and rifampicin synergy. A) Volcano plots of perfect guide gene-level knockdowns for colistin (0.44 µg/mL) + inducer (1mM IPTG) on the left and rifampicin (0.34 µg/mL) + inducer on the right, plotted by the significance (Stouffer’s *p*) and fitness (median log 2-fold change) compared to inducer only at T2. Horizontal dotted line denotes Stouffer’s *p* = 0.05, and vertical dotted lines denote fitness = ±1. Genes involved in NDH-1 and LOS biosynthesis are labeled. B) Heatmap with clustered dendrogram (k-means = 2) of relative fitness of NDH-1 and LOS genes in LB with inducer (1 mM IPTG) + colistin (0.44 µg/mL) or inducer + rifampicin (0.34 µg/mL) at T2 compared to T0. C) Knockdown-response curves for *lpxC* and *nuoB* in LB with inducer (1 mM IPTG), inducer + colistin (0.44 µg/mL), or inducer + rifampicin (0.34 µg/mL) at T1 and T2. Curves were fit to the fitness (log2-fold changes) of mismatch guides with a range of predicted knockdown for each gene.

NDH-1 pumps protons across the inner membrane to generate the proton motive force (PMF) that is required for ATP synthesis. Recent work has suggested that colistin can also disrupt the inner membrane by binding to LOS molecules that have yet to be trafficked to the outer membrane (45); thus, perturbation of the inner membrane by colistin may disrupt PMF formation by NDH-1. To validate or colistin results, we confirmed the resistant phenotype of *lpxC* and the sensitivity phenotype of *nuoB* using knockdown response curves of mismatch guides (Fig. 4C) and growth curves of individual strains (Fig. S7B). Because NDH-1 function is tightly coupled to fitness in *A. baumannii*, we suggest that indirect inhibition of NDH-1 is a key mechanism for growth inhibition by colistin.

Rifampicin is a relatively large antibiotic (822.9 Da) that targets RNA polymerase (RNAP) in the cytoplasm but is typically avoided for treating Gram-negative infections due to low permeability (46). Consistent with a permeability barrier to rifampicin function (47), we found that knockdown of LOS synthesis and transport genes strongly sensitized cells to rifampicin (FDR = 7.36e^-06^, AFC test). Unexpectedly, maximal knockdown of genes encoding NDH-1 caused cells to become resistant to rifampicin by an unknown mechanism, possibly by perturbing PMF-dependent import (FDR = 2.76e^-06^, AFC test; (Fig. 4A and 4B)). We validated the rifampicin sensitivity phenotype of *lpxC* and the resistance phenotype of *nuoB* using knockdown response curves (Fig. 4C) and growth curves of individual *lpxC* and *nuoB* knockdowns (Fig. S7).

Colistin and rifampicin showed strong, anticorrelated phenotypes (Fig. 4B), with LOS knockdowns causing resistance to colistin and sensitivity to rifampicin, and NDH-1 knockdowns resulting in sensitivity to colistin and resistance to rifampicin; these results support a collateral sensitivity mechanism for synergy (Fig. 5) (48). Indeed, knockdown of LOS or NDH-1 altered, but did not eliminate colistin-rifampicin synergy (Fig. S7A). Taken together, colistin and rifampicin synergistically kill *A. baumannii* by exerting opposite effects on LOS and NDH-1 that cannot be compensated for by reducing the function of either pathway.

**Fig. 5.**
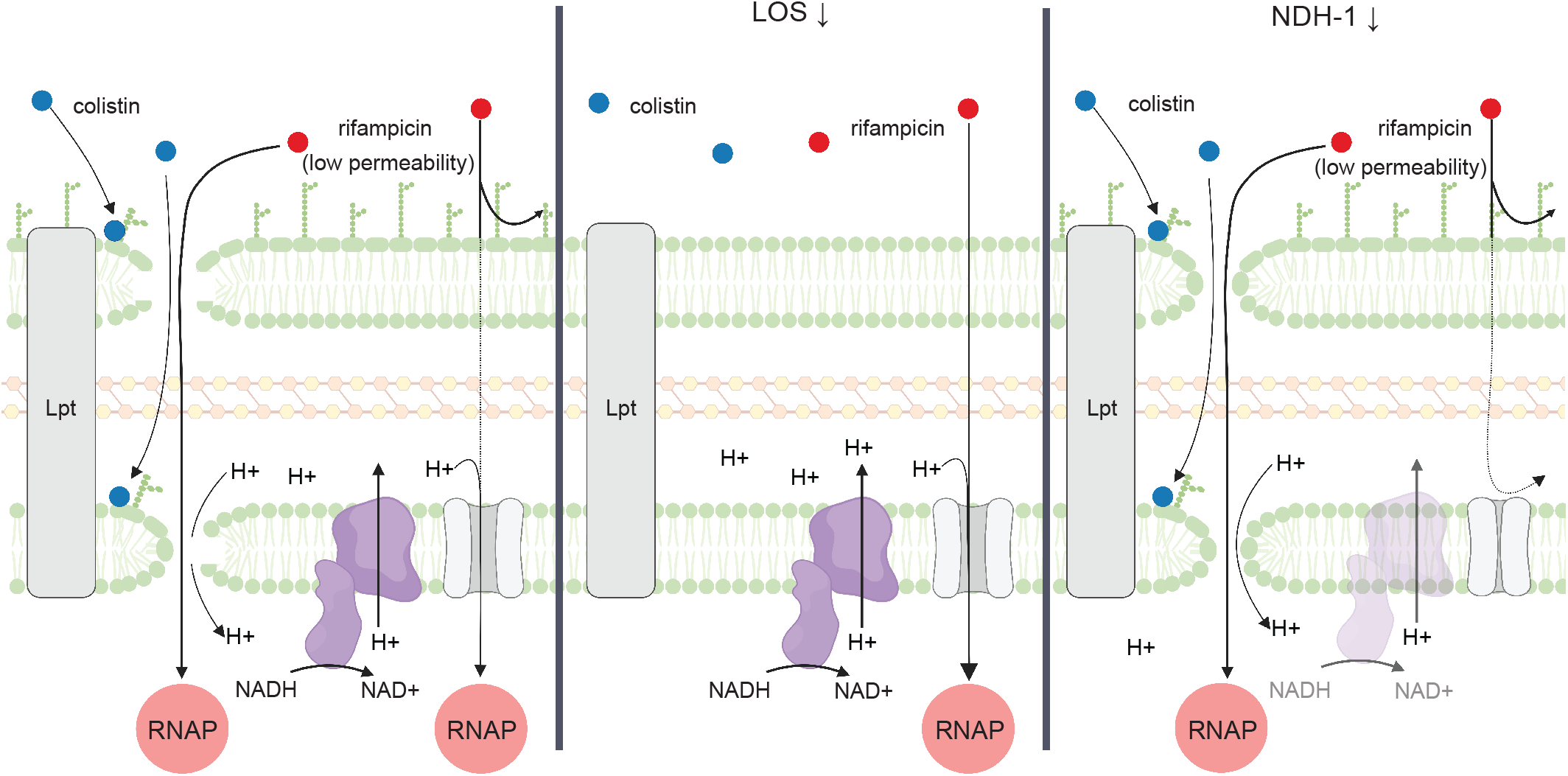
Model of colistin-rifampicin mechanism of synergy. Colistin disrupts membranes by binding to LOS, increasing permeability to rifampicin. Reduced expression of LOS also increases rifampicin permeability, but increases resistance to colistin. Reduced expression of NDH-1 disrupts proton motive force (PMF), increasing sensitivity to colistin while conferring resistance to rifampicin, possibly by reducing uptake through PMF-dependent transporters.

## Discussion

Bacterial susceptibility to antibiotics is underpinned by species- and condition-specific gene essentiality. The lack of innovative treatments for *A. baumannii* and other Gram-negative pathogens is directly attributable to our limited knowledge of genetic weaknesses in these bacteria. This work advances our understanding of genetic vulnerabilities in *A. baumannii* by systematically perturbing and phenotyping essential genes. Using CRISPRi to deplete essential gene products, we identified genes that are vulnerable to depletion as well as genes that synergize with or antagonize antibiotic action. Together, these studies define high-value targets for antibiotic discovery and provide a genetic rationale for synergistic therapies that may be broadly applicable.

Our study of essential gene depletion phenotypes in *A. baumannii* points to both unique and shared vulnerabilities with other bacterial species. Our finding that *A. baumannii* is highly sensitive to depletion of genes encoding NDH-1 (*nuo*) highlights a unique weakness in pathogens that are obligate aerobes and a possible therapeutic target. Among the Gram-negative ESKAPE pathogens (*Klebsiella, Acinetobacter, Pseudomonas*, and *Enterobacter*), only *A. baumannii* is known to require *nuo* genes for aerobic growth in rich medium (49). Recent from Manoil and colleagues in the non-pathogenic model strain, *Acinetobacter baylyi*, found that genes involved in oxidative phosphorylation were among the first to be depleted from a pool of transposon mutants (16); combining these observations with our CRISPRi results suggests that oxygen-dependent energy production is a physiological linchpin across the *Acinetobacter* genus. Our finding that *A. baumannii* genes involved in PG synthesis and translation are vulnerable to depletion underscores the conserved importance of these pathways across bacterial species (20, 33) and their foundational role as antibiotic targets. Our data suggest that NDH-1 inhibition plays a key role in colistin mode of action against *A. baumannii*. Edwards and colleagues (45) recently proposed a mechanism in which colistin kills bacteria by binding to LPS molecules in the inner membrane that have yet to be trafficked to the outer membrane by the Lpt complex; at high colistin concentrations, the inner membrane would be disrupted leading to cell lysis. However, at low colistin concentrations, we predict that perturbation of the inner membrane would mostly result in disruption of the proton gradient established by NDH-1 activity (Fig. 5). Thus, knockdown of genes encoding NDH-1 and treatment with colistin synergize to disrupt the proton gradient, leading to lethality. Consistent with our hypothesis, the proton gradient uncoupler CCCP synergizes with colistin in killing *A. baumannii* by a mechanism independent of efflux pump activity or expression (50). Although we prefer an indirect inhibition model, there is evidence that colistin can inhibit type II NADH dehydrogenase activity in a purified system at concentrations ∼100X higher than used in our experiments (51); it is unclear if this inhibition has physiological or medical relevance. In contrast, concentrations of colistin that are sublethal to *A. baumannii* as a monotherapy could be clinically relevant in the context of combination therapy, particularly since toxicity is a major dose-limiting concern of polymyxin antibiotics (52). Employing effective combination treatments using colistin concentrations below toxicity thresholds would greatly improve its clinical utility and safety against *A. baumannii*. Our CRISPRi approach to combination therapy screening could inform not only combinations with polymyxin antibiotics, but also other antibiotics which have dose-limiting toxicity concerns that prevent more widespread use.

Colistin and rifampicin have been shown to synergistically kill *A. baumannii* and other Gram-negatives (53), but the mechanism of synergy has remained elusive. Here, we propose a collateral sensitivity mechanism (48), in which genetic perturbations that promote colistin resistance increase sensitivity to rifampicin and vice versa (Fig. 5). Treatment with colistin selects for mutations in LOS biosynthesis genes (9), while the loss of LOS promotes permeability to rifampicin (and other antibiotics (53)). Accordingly, the presence of rifampicin has been shown to reduce recovery of inactivated *lpx* genes in colistin selections (54).

Mutations in *nuo* genes are commonly obtained in screens for tobramycin resistance in *P. aeruginosa* (55, 56) supporting a model in which reduced NDH-1 function decreases permeability of the inner membrane to antibiotics. Although the permeability mechanism is unknown, it is assumed that PMF-dependent transporters are responsible for tobramycin entry into the cytoplasm. By extension, we speculate that rifampicin traverses the inner membrane via one or more PMF-dependent transporters. Collateral sensitivity comes into play when mutations that reduce NDH-1 activity to block rifampicin entry increase sensitivity to colistin which further reduces NDH-1 function. This mechanism of synergy can impact other Gram-negative ESKAPE pathogens but is particularly relevant in *A. baumannii* because LOS is not essential, and NDH-1 is uniquely required for viability. In the case of colistin and rifampicin, collateral sensitivity manifests as anticorrelated phenotypes in chemical genomics data. We speculate that anticorrelated phenotypic signatures are predictive of antibiotic synergy, particularly in the context of bacteria with low permeability such as *A. baumannii* and *P. aeruginosa*. Interrogating a larger chemical genomics dataset with a greater diversity of antibiotics for these organisms will shed light on general rules for antibiotic-gene interactions and their implications for discovering synergy.

## Materials and Methods

### Strains and growth conditions

Strains are listed in Table S1. Details of strain growth conditions are described in the Supporting Information.

### General molecular biology techniques and plasmid construction

Plasmids and construction details are listed in Table S2. Oligonucleotides are listed in Table S3. Details of molecular biology techniques are described in the Supporting Information.

### *A. baumannii* Mobile-CRISPRi system construction

An *A. baumannii* strain with the Mobile-CRISPRi (MCi) system from pJMP1183 (21) inserted into the attTn7 site (Fig. S1A), which constitutively expresses mRFP and has an mRFP-targeting sgRNA, has a growth defect when induced with 1mM IPTG (Fig. S1B; “parent”). Strains with suppressors of the growth defect that still maintained a functional CRISPRi system were identified by plating on LB supplemented with 1mM IPTG and selecting white colonies (red colonies would indicate a no longer functional MCi system; Fig. S1B and S1C). gDNA was extracted and mutations in the dCas9 promoter were identified by Sanger sequencing (Fig. S1D). The Mobile-CRISPRi plasmid pJMP2748 is a variant of pJMP2754 (Addgene 160666) with the sgRNA promoter derived from pJMP2367 (Addgene 160076) and the dCas9 promoter region amplified from the *A. baumannii* suppressor strain gDNA with oJMP635 and oJMP636. Plasmid pJMP2776, which was used to construct the *A. baumannii* essential gene library and individual sgRNA constructs, was created by removal of the GFP expression cassette from pJMP2748 by digestion with PmeI and re-ligation. This system shows ∼20-fold knockdown when targeting the *GFP* gene (Fig. S1E). Plasmids will be submitted to Addgene.

### *A. baumannii* Mobile-CRISPRi individual gene and gene library construction

sgRNAs were designed to knockdown essential genes in *A. baumannii* 19606 using a custom python script and Genbank accession #s CP046654.1 and CP046655.1 as detailed in reference (19). sgRNA-encoding sequences were cloned between the BsaI sites of Mobile-CRISPRi (MCi) plasmid pJMP2776. Methodology for cloning individual guides was described previously in detail (19). Briefly, two 24-nucleotide (nt) oligonucleotides encoding an sgRNA were designed to overlap such that when annealed, their ends would be complementary to the BsaI-cut ends on the vector.

The pooled essential gene CRISPRi library was constructed by amplification of sgRNA-encoding spacer sequences from a pooled oligonucleotide library followed by ligation into the BsaI-digested MCi plasmid. Specifically, a pool of sgRNA-encoding inserts was generated by PCR amplification with primers oJMP697 and oJMP698 from a 78-nt custom oligonucleotide library (2020-OL-J, Agilent) with the following conditions per 500 µl reaction: 100 µl Q5 buffer, 15 µl GC enhancer, 10 µl 10mM each dNTPs, 25 µl each 10 µM primers oJMP897 and oJMP898, 10 µl 10 nM oligonucleotide library, 5 µl Q5 DNA polymerase, and 310 µl H2O with the following thermocycling parameters: 98°C, 30s; 15 cycles of: 98°C, 15s; 56°C, 15s; 72°C, 15s; 72°C, 10 min; 10°C, hold. Spin-purified PCR products were digested with BsaI-HF-v2 (R3733; NEB) and the size and integrity of full length and digested PCR products were confirmed on a 4% agarose e-gel (Thermo). The BsaI-digested PCR product (without further purification) was ligated into a BsaI-digested MCi plasmid as detailed in (19). The ligation was purified by spot dialysis on a nitrocellulose filter (Millipore VSWP02500) against 0.1 mM Tris, pH 8 buffer for 20 min prior to transformation by electroporation into *E. coli* strain BW25141 (sJMP3053). Cells were plated at a density of ∼50,000 cells/plate on 150mm LB-2% agar plates supplemented with carbenicillin. After incubation for 14 h at 37°C, colonies (∼900,000 total) were scraped from the agar plates into LB, pooled, and the plasmid DNA was extracted from ∼1×10^11^ cells using a midiprep kit. This pooled Mobile-CRISPRi library was transformed by electroporation into *E. coli* mating strain sJMP3049, plated at a density of ∼50,000 cells/plate on 150mm LB-2% agar plates supplemented with carbenicillin and DAP. After incubation for 14 h at 37°C, colonies (∼1,000,000 total) were scraped from the agar plates and pooled, the OD600 was normalized to 27 in LB with DAP and 15% glycerol and aliquots of the pooled CRISPRi library were stored as strain sJMP2942 at -80°C.

### Transfer of the Mobile-CRISPRi system to the *A. baumannii* chromosome

The MCi system was transferred to the Tn*7att* site on the chromosome of *A. baumannii* by quad-parental conjugation of three donor strains—one with a mobilizable plasmid (pTn7C1) encoding Tn7 transposase, another with a conjugal helper plasmid (pEVS104), and a third with a mobilizable plasmid containing a Tn7 transposon encoding the CRISPRi system—and the recipient strain *A. baumannii* 19606. A detailed mating protocol for strains with individual sgRNAs was described previously (19). Briefly, 100 µl of culture of donor and recipient strains were added to 600 µl LB, pelleted at ∼8000 x g, washed twice with LB prior to depositing cells on a nitrocellulose filter (Millipore HAWP02500) on an LB plate, and incubated at 37°C, ∼5 hr. Cells were removed from the filter by vortexing in 200 µl LB, serially diluted, and grown with selection on LB-gent plates at 37°C.

For pooled library construction, Tn*7* transposase donor (sJMP2644), conjugation helper strain (sJMP2935), and recipient strain (sJMP490) were scraped from LB plates with appropriate selective additives into LB and the OD600 was normalized to ∼9. An aliquot of sJMP2942 pooled library strain was thawed and diluted to OD600 of ∼9. Eight ml of each strain was mixed and centrifuged at 8000xg, 10 min. Pelleted cells were resuspended in 4 ml LB, spread on two LB agar plates, and incubated for 5hr at 37°C prior to resuspension in LB + 15% glycerol and storage at -80°C. Aliquots were thawed and serial dilutions were plated on LB supplemented with gent (150) and LB. Efficiency of trans-conjugation (colony forming units on LB-gent vs. LB) was ∼1 in 10^7^. The remaining frozen stocks were plated on 150 mm LB plates solidified with 2% agar and supplemented with gent (150) and incubated for 16 h at 37°C. Cells were scraped from plates and resuspended in EZRDM (Teknova) + 25mM succinate + 15% glycerol at OD600 = 15 and aliquots were stored at -80°C as strain sJMP2949.

### Library growth experiment

The *A. baumannii* essential gene CRISPRi library (sJMP2949) was revived by dilution of 50 µl frozen stock (OD600 = 15) in 50 ml LB (starting OD600 = 0.015) and incubation in 250 ml flasks shaking at 37°C until OD600 = 0.2 (∼2.5 h) (timepoint = T0). This culture was diluted to OD600 = 0.02 in 4 ml LB with 1mM IPTG and antibiotics (colistin, imipenem, meropenem, rifampicin, and no antibiotic control) in 14 ml snap cap culture tubes (Corning 352059) in duplicate and incubated with shaking for 18 h at 37°C (T1). These cultures were serially diluted back to OD600 = 0.01 into fresh tubes containing the same media and incubated with shaking for 18 h at 37°C again (T2) for a total of ∼10-15 doublings. Cells were pelleted from 1 ml of culture in duplicate at each time point (T0, T1, T2) and stored at -20°C. Final antibiotic concentrations were (in µg/ml): colistin (Sigma C4461): 0.44 and 0.67, imipenem (Sigma I0160): 0.06 and 0.09, meropenem (Sigma 1392454): 0.11 and 0.17, and rifampicin (Sigma R3501): 0.34.

### Sequencing library samples

DNA was extracted from cell pellets with the DNeasy gDNA extraction kit (Qiagen) according to the manufacturer’s protocol, resuspending in a final volume of 100 µl with an average yield of ∼50 ng/µl. The sgRNA-encoding region was amplified using Q5 DNA polymerase (NEB) in a 100 µl reaction with 2 µl gDNA (∼100 ng) and primers oJMP697 and oJMP698 (nested primers with adapters for index PCR with Illumina TruSeq adapter) according to the manufacturer’s protocol using a BioRad C1000 thermalcycler with the following program: 98°C, 30s then 16 cycles of: 98°C, 15s; 65°C, 15s; 72°C, 15s. PCR products were purified using the Monarch PCR and DNA Cleanup and eluted in a final volume of 20 µl for a final concentration of ∼20 ng/µl).

Samples were sequenced by the UW-Madison Biotech Center Next Generation Sequencing Core facility. Briefly, PCR products were amplified with nested primers containing i5 and i7 indexes and Illumina TruSeq adapters followed by bead cleanup, quantification, pooling and running on a Novaseq 6000 (150bp paired end reads).

### Library data analysis

For more information on digital resources and links to custom scripts, see Table S4.

### Counting sgRNA Sequences

Guides were counted using *seal*.*sh* script from the *bbtools* package (Release: March 28, 2018). Briefly, paired FASTQ files from amplicon sequencing were aligned in parallel to a reference file corresponding to the guides cloned into the library. Alignment was performed using *k*-mers of 20 nucleotide length—equal to the length of the guide sequence.

### Condition Comparisons – Quantification and Confidence

Log2-fold change and confidence intervals were computed using *edgeR*. Briefly, trended dispersion of guides was estimated and imputed into a quasi-likelihood negative binomial log-linear model. Changes in abundance and the corresponding false discovery rates were identified for each guide in each condition individually. Finally, log2-fold abundance changes were calculated by taking the median guide-level log2-fold change; confidence was calculated by computing the Stouffer’s *p*-value (*poolr R* package) using FDR for individual guides across genes.

### Knockdown-Response Curves

Code was adapted from the *drc* (*DoseResponse*) *R* package to generate 4-parameter logistic curves describing the relationship between predicted knockdown (independent) and the log2-fold change in strain representation (dependent) for all (∼10) mismatch guides per gene.

## Data Sharing Plan

Raw data will be deposited in the Sequence Read Archive (SRA), code used to analyze the data will be available on Github, and plasmids will be available from Addgene. Other reagents and protocols are available upon request.

## Supporting information

Supplemental Tables List

## Acknowledgements

This work was supported by a Career Transition Award from the NIH National Institute of Allergy and Infectious Diseases (K22AI137122). R.D.W. was supported by the Predoctoral Training Program in Genetics (NIH 5T32GM007133). J.S.T. was supported by the Biotechnology Training Program (NIH 5T32GM135066) and a GRFP from the NSF. We thank Agilent Technologies for providing SurePrint Oligonucleotide libraries and Laura Whitman for oligo synthesis support and the University of Wisconsin Biotechnology Center for technical support with Illumina sequencing.

