## Supplementary figures and images for "Essential Gene Phenotypes Reveal Antibiotic Mechanisms and Synergies in *Acinetobacter baumannii*"

### Supplemental Figure S1

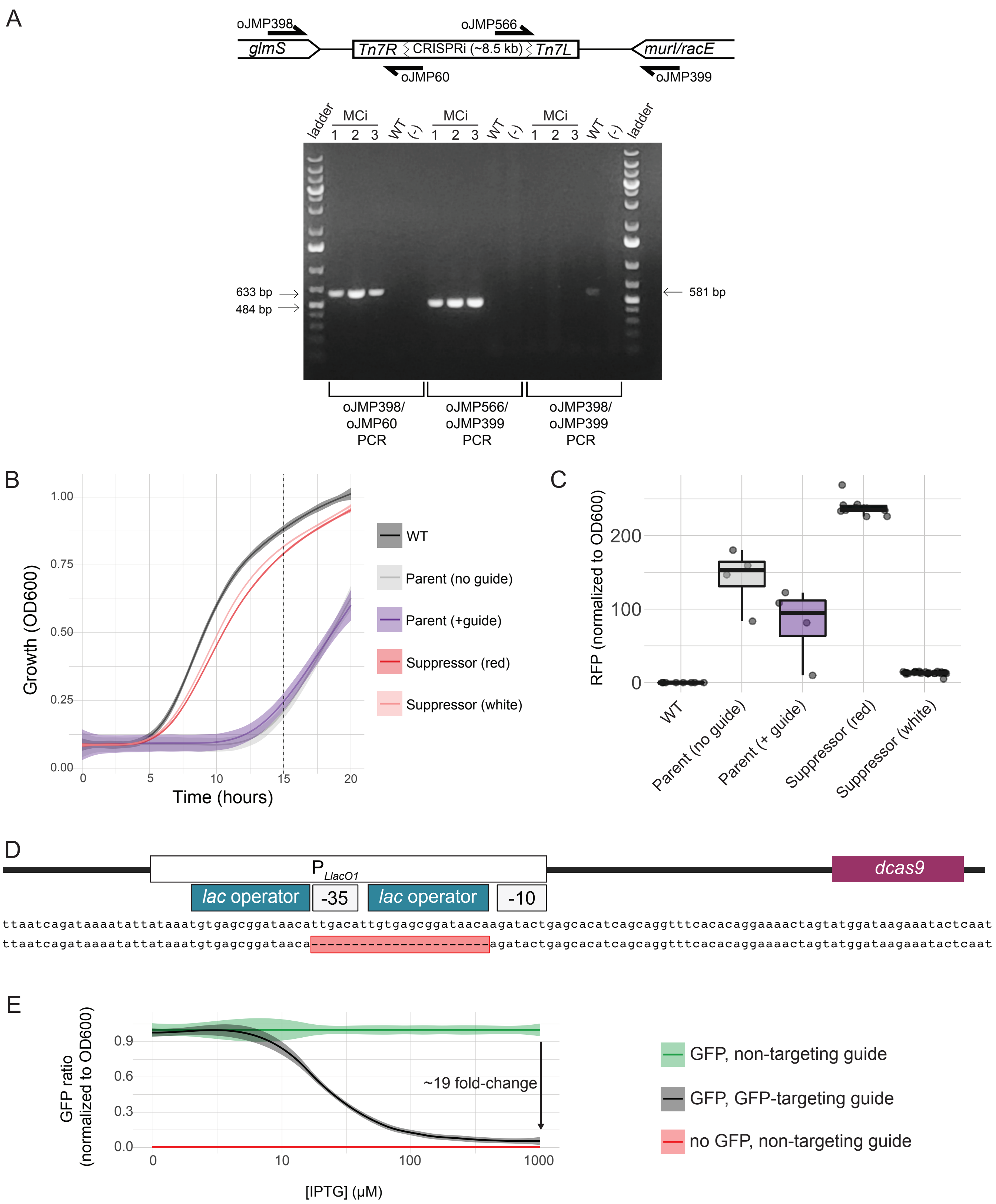

### Supplemental Figure S2

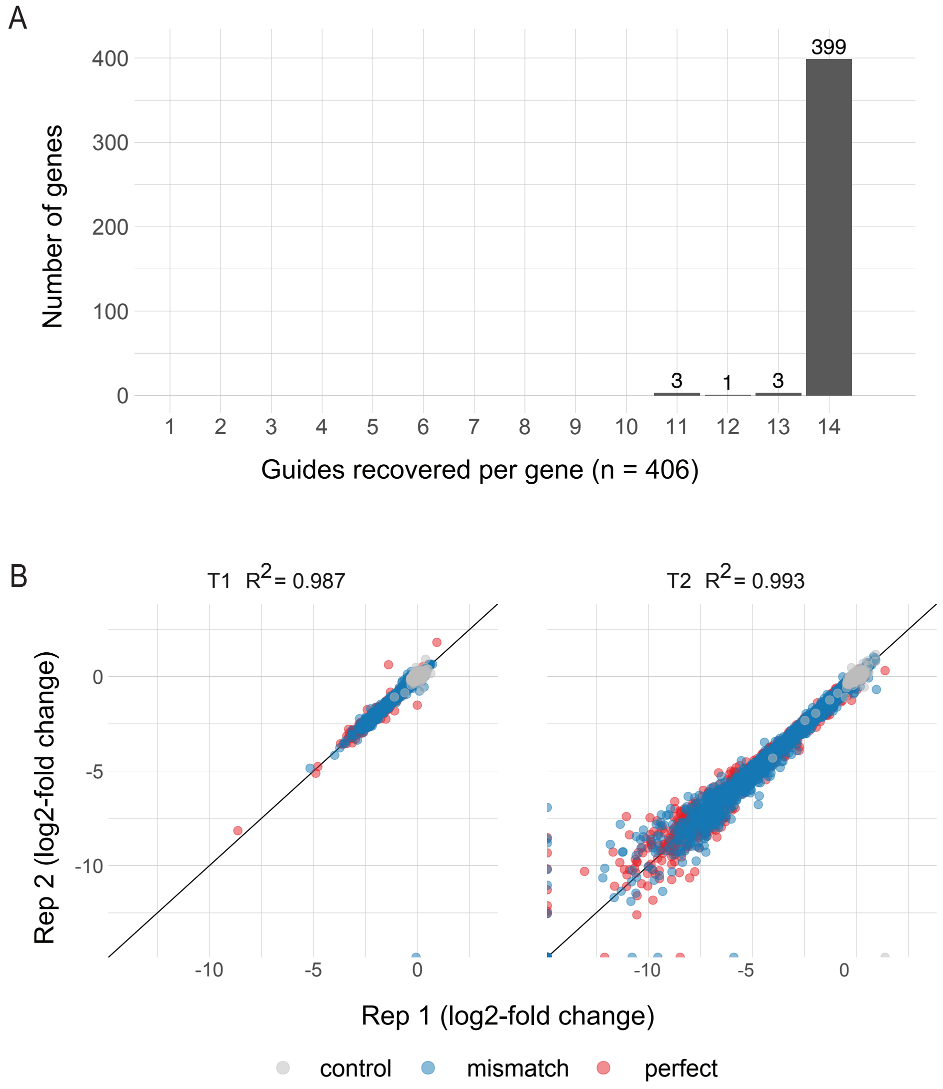

### Supplemental Figure S3

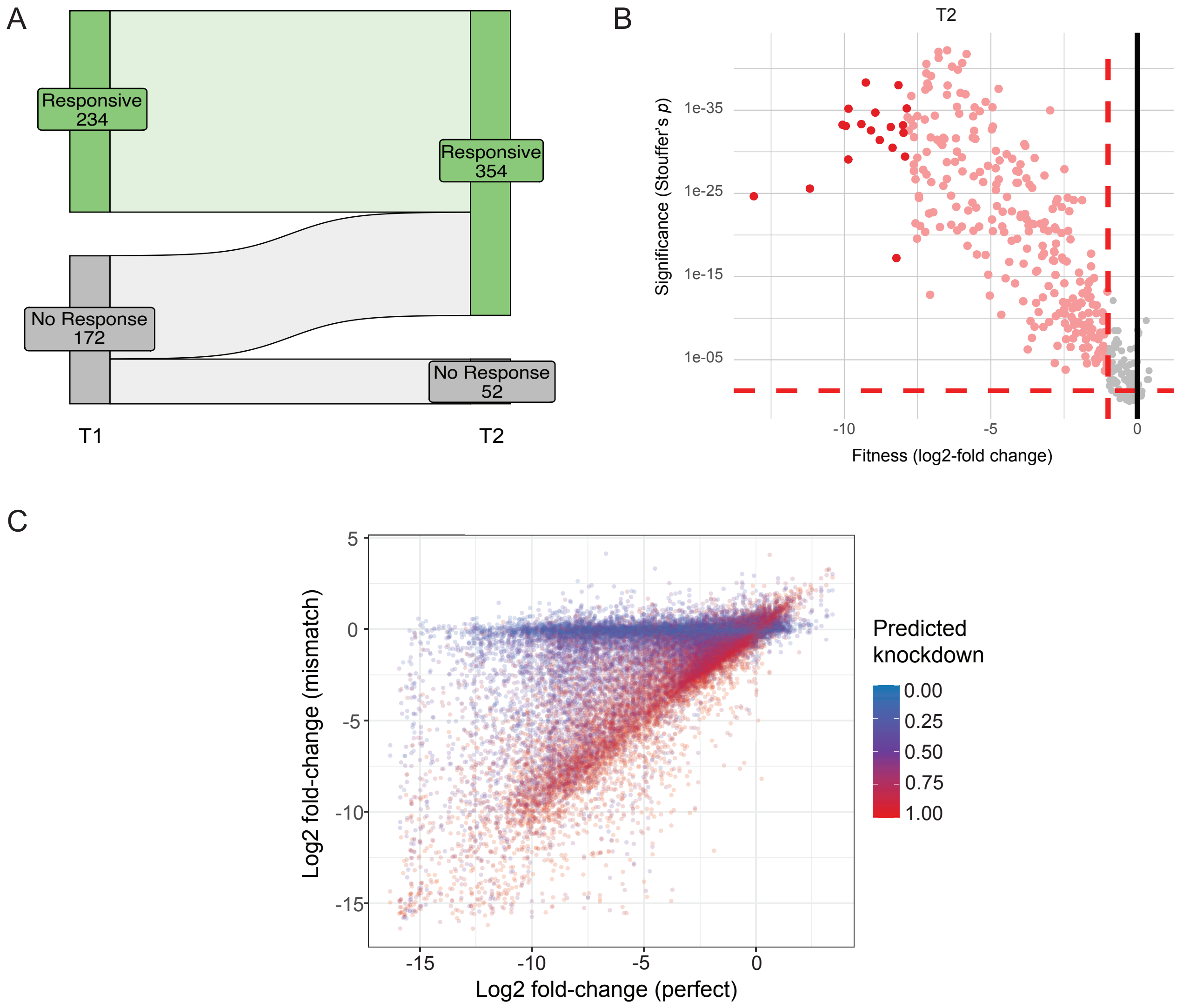

### Supplemental Figure S4

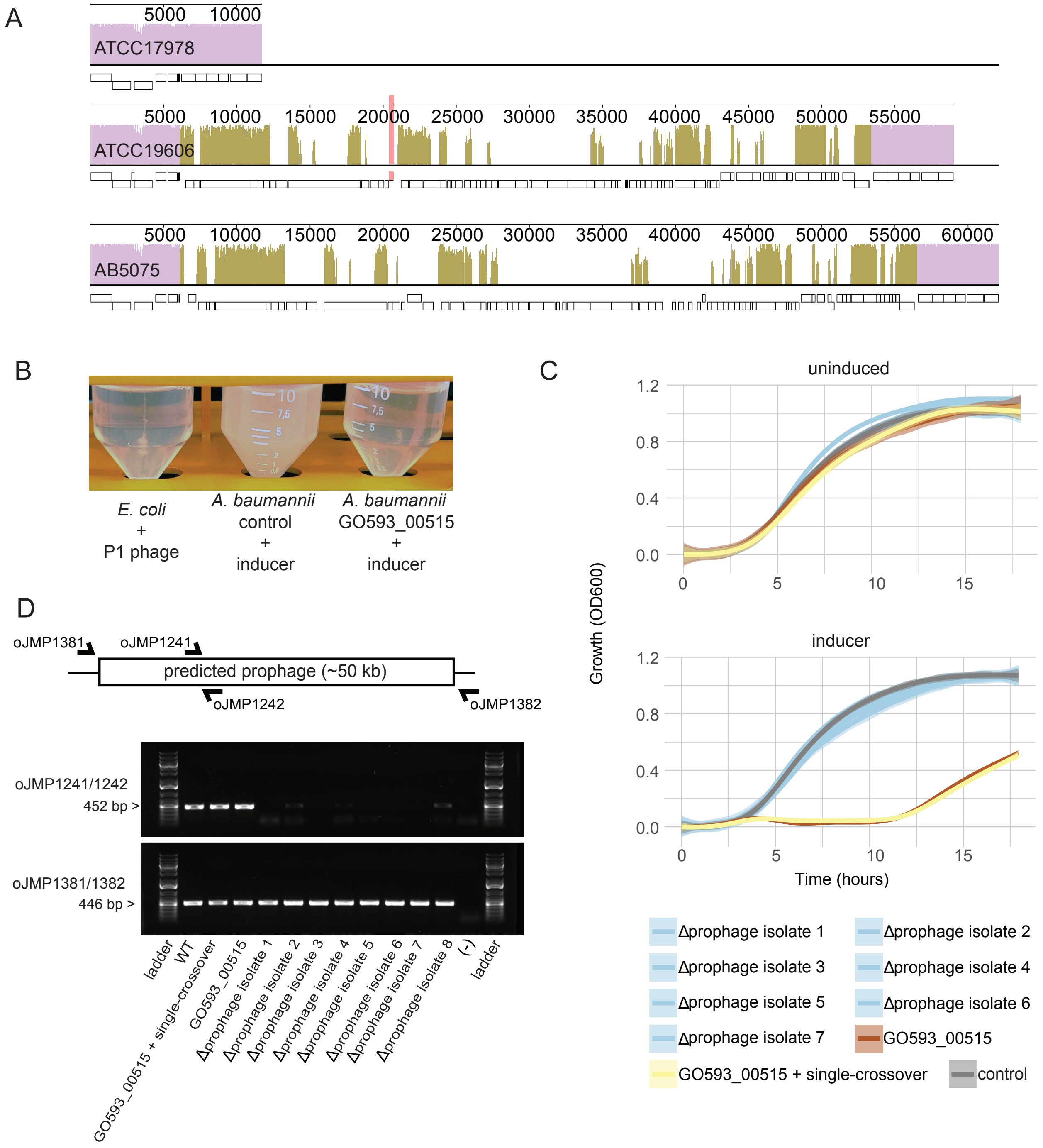

### Supplemental Figure S5

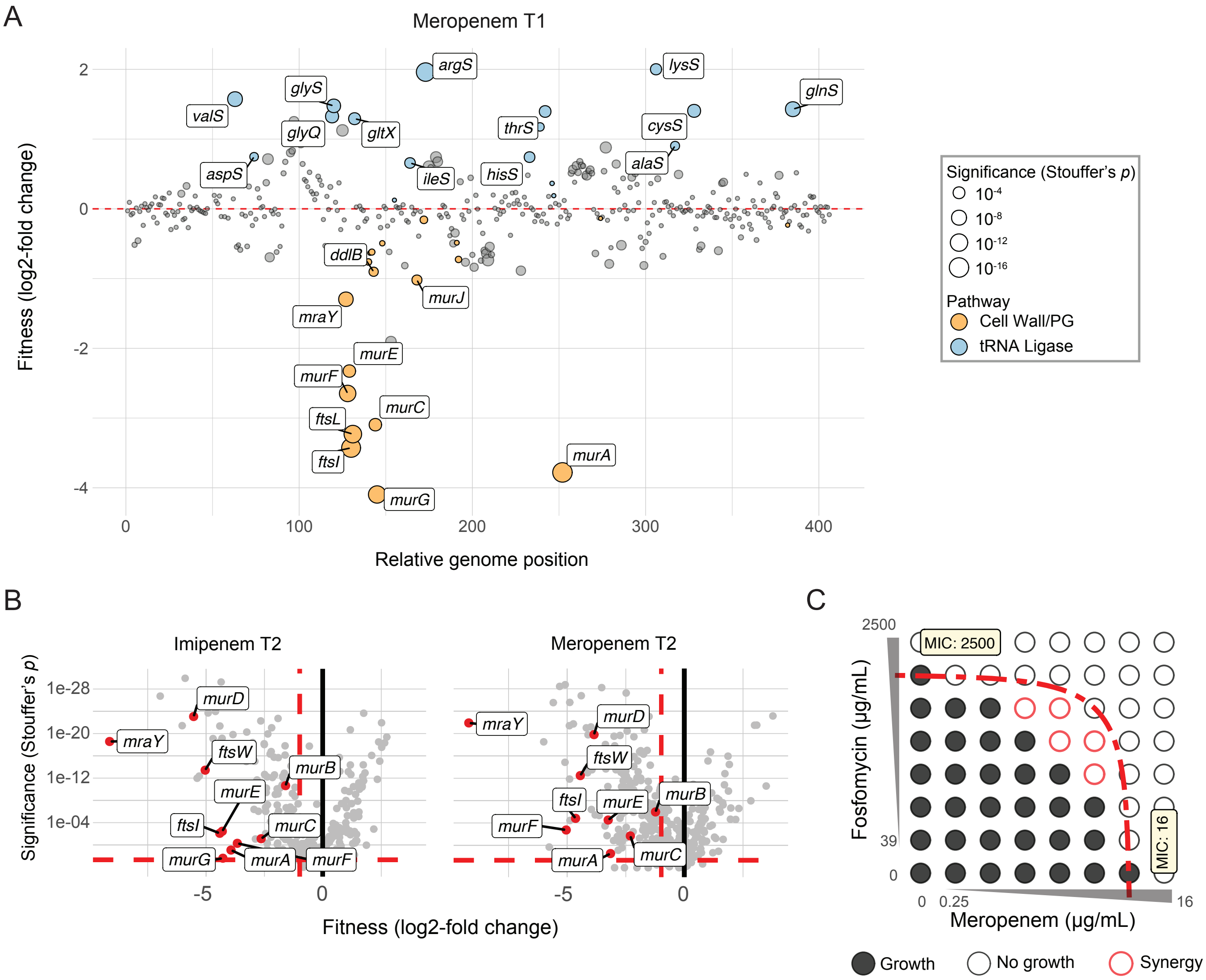

### Supplemental Figure S6

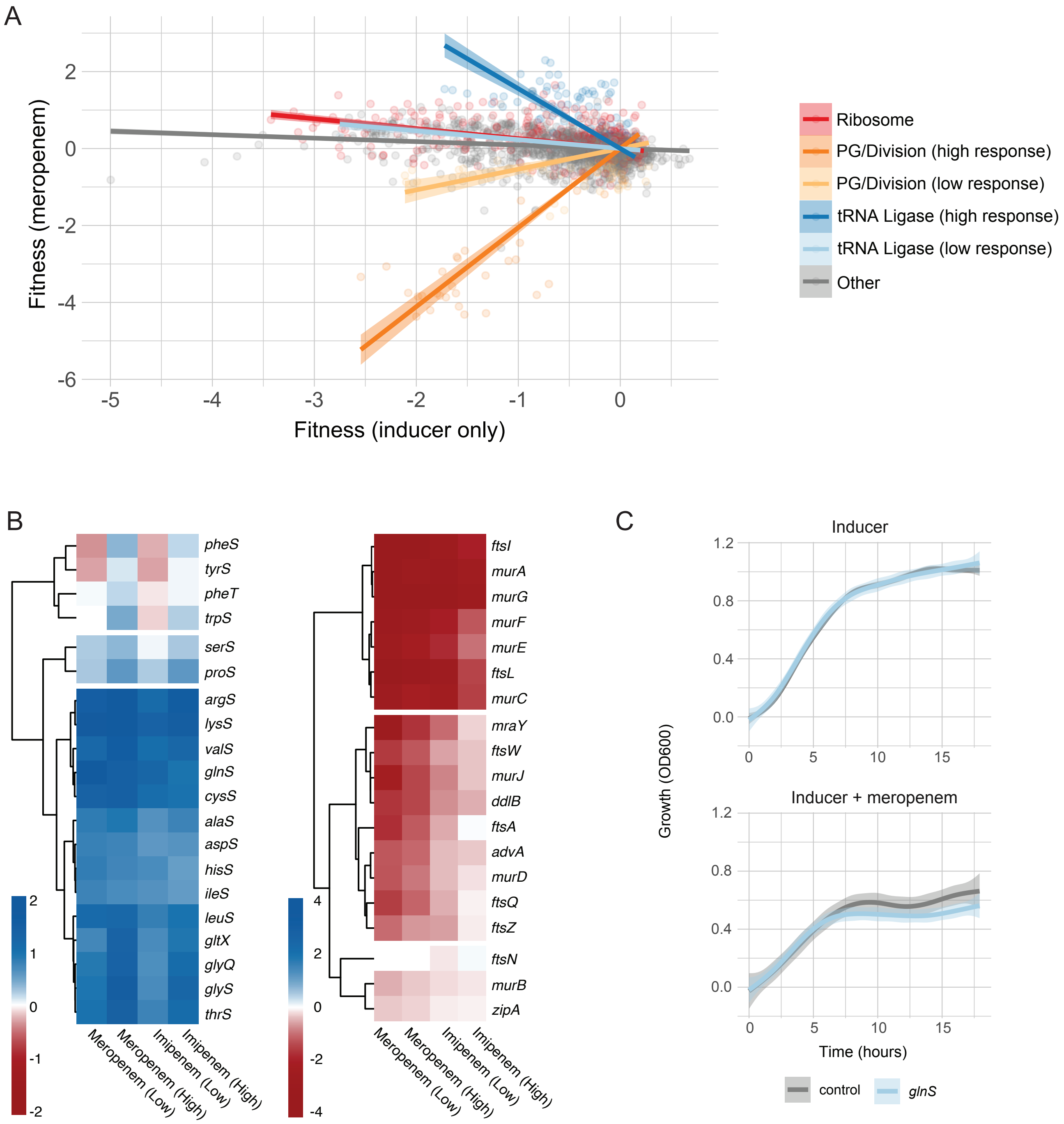

### Supplemental Figure S7

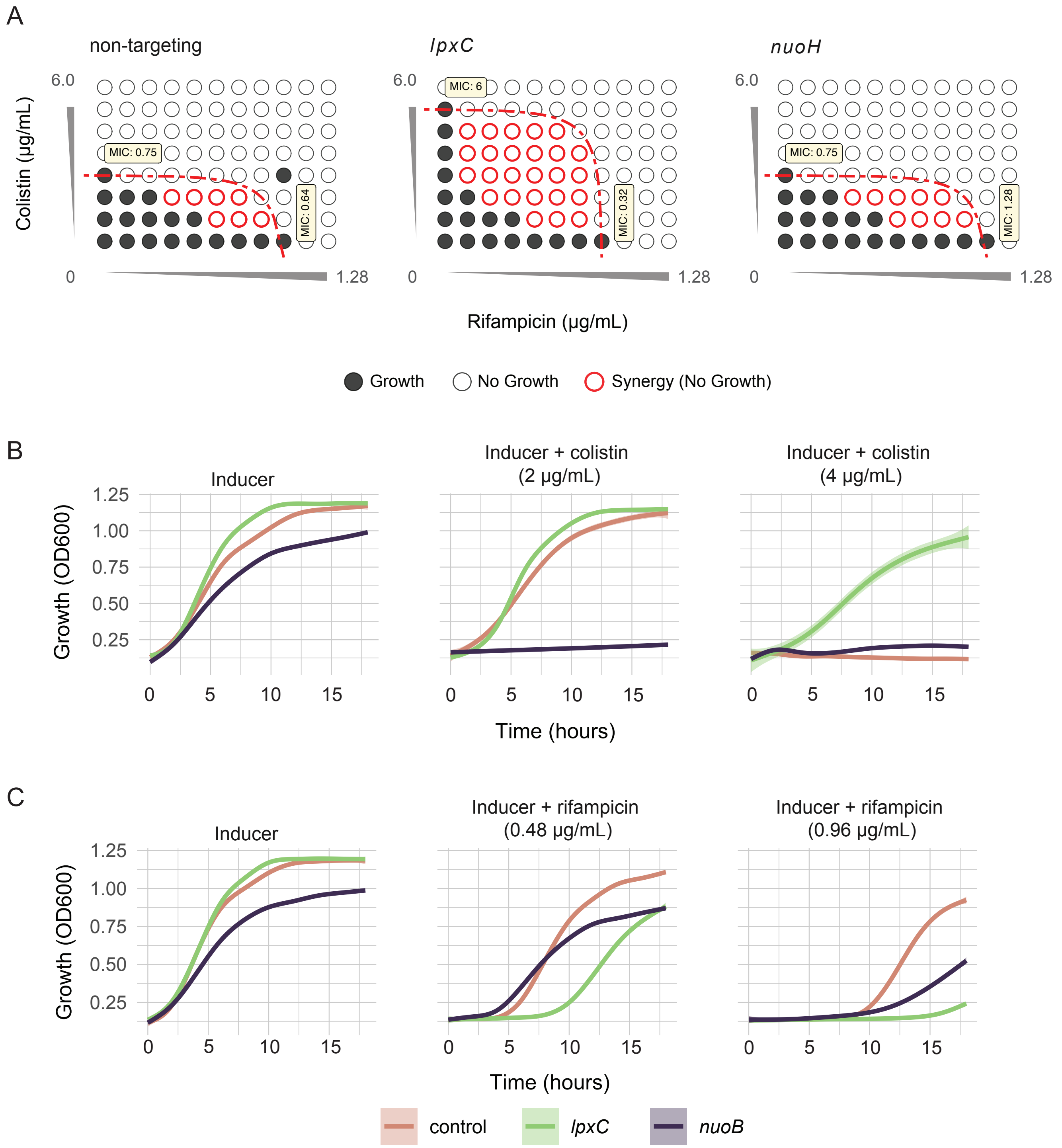
