## Supplemental Figure Legends for "Essential Gene Phenotypes Reveal Antibiotic Mechanisms and Synergies in *Acinetobacter baumannii*"

**Fig. S1. Construction and optimization of an *A. baumannii* CRISPRi library**

A) Agarose gel (1% agarose, 1X TAE buffer) electrophoresis of PCR products to confirm insertion of Mobile-CRISPRi downstream of glmS in *A. baumannii* using specified primer sets. GeneRuler 1 kb Plus ladder or template genomic DNA are denoted over each lane.
B) Growth, measured by OD600, over 20 hours of WT or CRISPRi constructs with RFP. Parent strain from [21]**.** IPTG-inducible CRISPRi system is toxic to cells when fully induced with 1mM IPTG, both without guide (no guide strain) and with guide (parent strain) for targeting RFP. Suppressors (both with *rfp*-targeting guide) show recovery of growth. White suppressor strain retains a functional CRISPRi system. Dotted line represents timepoint for RFP knockdown measurements in next panel.
C) Boxplot of OD600-normalized RFP measurements for CRISPRi constructs at 15 hours. Dots represent biological replicates. Suppressor (white) is a non-toxic variant that retains RFP knockdown.
D) Map of Mobile-CRISPRi dCas9 promoter showing original $P_{LlacO1}$ promoter sequence (top) and *A. baumannii* mutant promoter sequence (bottom).
E) Graph showing ratio of GFP remaining across IPTG concentrations compared to the mean of GFP, non-targeting guide control strain. Strain expressing GFP with GFP-targeting guide shows ~19 fold-change in normalized fluorescence with induction at 1mM IPTG. Strain with no GFP and non-targeting guide acts as negative control.s

**Fig. S2. Validation of *A. baumannii* CRISPRi library**

A) Histogram of guides recovered per gene in the library (n = 406).
B) Scatterplot comparing log2-fold changes of all guides in biological replicates for T1 and T2 samples after library induction. The solid line is y = x.

**Fig. S3. Identification of *A. baumannii* essential genes sensitive to deletion**

A) Sankey plot of genes at T1 and T2 in all conditions, binned as no response or responsive. Genes are responsive if gene-level knockdowns have log2-fold changes > 1 or <-1 and Stouffer's p < 0.05.
B) Volcano plot of gene-level knockdowns at T2, plotted by the significance (Stouffer's *p*) and fitness (median log2-fold change) across perfect guides. Gene-level knockdowns with log2-fold change < -1 and Stouffer's *p* < 0.05 are colored pink, and knockdowns with log2-fold change < -7.85 (top 5% in response) are colored red.
C) Scatterplot, where each dot represents the log2-fold changes of a perfect guide and a corresponding mismatch, colored for the predicted knockdown of that mismatch. Mismatch guides with low predicted knockdown show a different apparent trend than those with high predicted knockdown.

**Fig. S4. GO593_00515 is conditionally essential**

A) Mauve alignment (see Table S4) of *A. baumannii* 19606 genomic locus containing GO593_00515 (pink) and surrounding prophage (yellow) to 17978 and AB5075. Regions of same color represent alignment between genomes.
B) Cultures seven hours after addition of P1 phage lysate to *E. coli* MG1655 (left) and addition of inducer (1 mM IPTG) to *A. baumannii* non-targeting control strain (middle) or GO593_00515 knockdown strain (right).
C) Growth, measured by OD600, over 18 hours for a non-targeting control strain, GO593_00515 knockdown strain, GO593_00515 knockdown strain with single-crossover of prophage deletion plasmid (intermediate in construction), and multiple strains of GO593_00515 knockdown with deleted prophage in LB (uninduced, top) or LB with inducer (1 mM IPTG, bottom).
D) Agarose gel (1% agarose, 1X TAE buffer) electrophoresis of PCR products to confirm prophage deletion using specified primer sets. GeneRuler 1 kb Plus ladder or template genomic DNA are denoted below each lane. PCR products with oJMP1381/1382 for strains with prophage still present suggests prophage excision happens at some frequency in WT.

**Fig. S5. Carbapenem-gene interactions in *A. baumannii*.**

A) Bubble plots of fitness, or median log2-fold change, for each gene in meropenem (0.17 µg/mL) at T1 compared to the inducer only (no drug) sample. Bubbles are plotted by gene position, or relative location on the genome compared to other predicted essential genes. Significance (Stouffer's *p*) is represented by size; genes in PG/division (orange) or tRNA ligase (blue) pathways are highlighted.
B) Volcano plots of gene-level knockdowns at T2 for imipenem (0.09 µg/mL) and meropenem (0.17 µg/mL), plotted by the significance (Stouffer's p) and fitness (median log2-fold change) across perfect guides. Dotted lines represent log2-fold change = -1 and Stouffer's *p* = 0.05. Genes involved in PG/division are labeled.
C) Sensitivity assay for fosfomycin-meropenem interactions. 2-fold serial dilutions of drugs from MICs represented by gray wedges. No growth below dotted line (FIC index ≤ 0.5), shows synergy (red border).

**Fig. S6. Pathway responses to carbapenems in *A. baumannii.***

A) Scatterplot of mismatch guide fitness (log2-fold change) in inducer only compared to relative fitness in meropenem at T1. Lines represent linear model fits with 95% confidence interval. Guides for genes in PG/division or tRNA ligase pathways are divided into groups using hierarchical clustering based on response to meropenem as depicted in panel B.
B) Heat maps showing Ward method hierarchical clustering for responses of tRNA ligase (left) and PG/Division (right) gene groups. Genes with responses of greater magnitude cluster together (high response) while genes with responses of lower magnitude cluster separately (low response).
C) Growth, measured by OD600, over 18 hours for a non-targeting control strain and *glnS* knockdown strain in 1 mM IPTG (left) or 1 mM IPTG plus 0.17 µg/mL meropenem (right).

**Fig. S7. Genetic interactions with rifampicin and colistin in *A. baumannii*.**

A) Sensitivity assay for colistin-rifampicin interactions in nontargeting control strain (left), *lpxC* knockdown strain (middle), or *nuoH* knockdown strain (right). 2-fold serial dilutions of drugs from MICs represented by gray wedges. No growth below dotted line (FIC index ≤ 0.5) shows synergy (red border).
B) Growth, measured by OD600, over 18 hours for a non-targeting control strain, *lpxC* knockdown strain, and *nuoB* knockdown strain in LB with inducer (1 mM IPTG, left) or LB with inducer and colistin (middle, right).
C) Growth, measured by OD600, over 18 hours for a non-targeting control strain, *lpxC* knockdown strain, and *nuoB* knockdown strain in LB with inducer (1 mM IPTG, left) or LB with inducer and rifampicin (middle, right).
