## Supplemental Tables List for "Essential Gene Phenotypes Reveal Antibiotic Mechanisms and Synergies in *Acinetobacter baumannii*"

**Aba_lib manuscript Supplemental Tables list**

Supplemental Table 1: Strains

Supplemental Table 2: Plasmids

Supplemental Table 3: Oligonucleotides and Synthetic DNA

Supplemental Table 4: Electronic Resources

Supplemental Table 5: Oligo Library Sequences and Key

Supplemental Table 6: sgRNA Fitness Results

Supplemental Table 7: Gene Fitness Results

Supplemental Table 8: Ortholog Analysis and Annotation
