## Supplemental Methods for "Essential Gene Phenotypes Reveal Antibiotic Mechanisms and Synergies in *Acinetobacter baumannii*"

**SI Materials and Methods**

**Strains and growth conditions.** Strains are listed in Table S1. *Escherichia coli* and *Acinetobacter baumannii* were grown in Lennox lysogeny broth (LB) (10 g tryptone, 5 g yeast extract, 5 g NaCl per liter; BD 240230) at 37°C in a flask with shaking at 250 rpm, in a culture tube on a roller drum at max speed, in a 96 deep well plate with shaking at 900 rpm, or in a plate reader (Tecan Infinite 200 Pro Mplex) with shaking. Culture medium was solidified with 1.5%-2% agar for growth on plates. Antibiotics were added when necessary: for *E. coli*, 100 µg/ml ampicillin (amp) or carbenicillin (carb), 15 µg/ml gentamicin (gent), or 30 µg/ml kanamycin (kan); and for *A. baumannii*, 150 µg/ml gentamicin (gent) or 60 µg/ml kanamycin (kan). Diaminopimelic acid (DAP) was added at 300 µM to support growth of *E. coli* *dap*- donor strains. IPTG (isopropyl b-D-1-thiogalactopyranoside) (0 to 1 mM) was added where indicated in the figures or figure legends. Strains were preserved in 15% glycerol at -80°C. *pir*-dependent plasmids were propagated in *E. coli* strain BW25141 *att*Tn*7*::*acrIIA4* (sJMP3053) for DNA extraction and analysis or in *E. coli* strain WM6026 *att*Tn*7*::*acrIIA4* (sJMP3049) for conjugation.

**General molecular biology techniques and plasmid construction.** Plasmids and construction details are listed in Table S2. Oligonucleotides are listed in Table S3. Oligonucleotides were synthesized by Integrated DNA Technologies (Coralville, IA) except the pooled oligonucleotide libraries which were synthesized by Agilent (Santa Clara, CA). Plasmid DNA was purified using the GeneJet plasmid miniprep kit (K0503; Thermo Scientific) or the PureLink HiPure Plasmid Midiprep kit (K210005; Invitrogen). Genomic DNA was purified using the GeneJet genomic DNA purification kit (K0721; Thermo Scientific). DNA fragments were amplified by PCR with Q5 DNA polymerase (M0491; New England Biolabs (NEB)) or OneTaq DNA Polymerase (NEB). DNA was digested with restriction enzymes from NEB. DNA fragments were spin purified using the Monarch PCR & DNA cleanup kit (T1030; NEB) or the DNA Clean & Concentrator kit (D4004; Zymo Research) after digestion or amplification. Linearized plasmids were re-circularized using T4 DNA ligase (M0202; NEB). Plasmids were assembled from restriction enzyme linearized or PCR-amplified vector and PCR products or synthetic DNA fragments using the NEBuilder Hifi DNA assembly kit (E2621; NEB). Individual spacers were cloned into BsaI-digested Mobile-CRISPRi (MCi) plasmids by annealing of two complementary oligonucleotides followed by ligation with T4 DNA ligase (NEB). For additional technical details about cloning individual spacers, see below and reference (1). For details about the pooled CRISPRi library construction, see below. Plasmids were transformed into electrocompetent *E. coli* cells using a Bio-Rad Gene Pulser Xcell on the EC1 setting. Sanger DNA sequencing was performed by Functional Biosciences (Madison, WI). Next Generation Sequencing was performed by the UW-Madison Biotechnology Center Next Generation Sequencing Core using an Illumina NovaSeq 6000.

***E. coli* strain construction.** *E. coli* *pir*+ cloning (sJMP3053) and mating (sJMP3049) strains expressing the *Listeria monocytogenes* prophage type II-A CRISPR-Cas9 inhibitor protein encoding gene *acrIIA4* (2) **were constructed to inhibit dCas9 during Mobile-CRISPRi library construction and conjugation. Additionally, the WT *recA* allele in *E. coli* strain sJMP3049 was replaced with *recA1* to decrease homologous recombination.**

**Allelic replacement of chromosomal *recA* with *recA1*.** An FRT-flanked chloramphenicol resistance (FRT-*cat*-FRT) cassette was inserted on the *E. coli* BW25141 chromosome between *mltB* and *srlA* (closely linked to the *recA1* allele) to facilitate allelic replacement*.* The FRT-*cat*-FRT cassette was amplified by PCR from plasmid pJMP1356 (3) using primers oJMP211 and oJMP212 (which also have 40 nt homology to the chromosomal insertion site), DpnI digested to destroy the plasmid, and inserted onto the chromosome of strain *E. coli* strain BW25141 (sJMP146) by λ-Red-mediated recombination using pSIM19 (pJMP38) encoding the λ-Red proteins, as previously described (4), resulting in strain sJMP601. The allele was transferred to the *E. coli* WM6026 strain background (sJMP424) by P1–*vir*-mediated transduction, as previously described (5), resulting in strain sJMP604. The *cat* gene was removed by transformation with plasmid pCP20, encoding a constitutively expressed FLP recombinase (pJMP3008), as previously described (6), resulting in strain sJMP624. Identity of the *recA1* allele was confirmed by PCR with flanking primers (oJMP201 and oJMP202) followed by sequencing.

***att*Tn*7*::*acrIIA4* gene insertion.** A plasmid (pJMP3018) bearing a Tn7 transposon encoding the *acrIIA4* gene under the control of a strong constitutive synthetic promoter and a chloramphenicol resistance cassette (FRT-*cat*-FRT) and a plasmid (pJMP442) encoding Tn*7* transposase were co-electroporated into *E. coli* strain BW25113 (sJMP006), using chloramphenicol to select for transposition into the *att*Tn*7* site, resulting in strain sJMP3030. The *att*Tn*7::acrIIA4* allele (Cam^r^) was transferred to *E. coli* strain BW25141 or WM6026 expressing *recA* from an unstable mini-F plasmid (sJMP345 or sJMP3040, respectively) by P1–*vir*-mediated transduction, as previously described (5). After serial passaging without selection, the resultant strains after plasmid loss are sJMP3034 or sJMP3043, respectively. The *cat* gene was removed by transformation with a plasmid encoding a constitutively expressed FLP recombinase (pJMP3008), as previously described (6) resulting in final strains sJMP3053 (*E. coli* strain BW25113 expressing *acrIIA4*) and sJMP3049 (*E. coli* strain WM6026 expressing *acrIIA4*).

***A. baumannii* Mobile-CRISPRi system characterization**

**Growth assay.** Growth of *A. baumannii* 19606 WT (sJMP490) and *att*Tn*7*::Mobile-CRISPRi (sJMP6335) in LB with and without 1mM IPTG were compared to assess whether the Mobile-CRISPRi system causes a growth defect. Four individual isolates were grown overnight in 300µl LB in a deep well microtiter plate with shaking at 900rpm. Cultures were diluted 1:1000 in LB or LB + 1mM IPTG and incubated in a plate reader for 16 hr at 37°C with shaking and growth was monitored by measuring OD600 every 15 min. This assay was repeated 3 times with 4 independent colonies. Growthcurver (7) was used to compare growth parameters.

**Induction assay.** Induction of the Mobile-CRISPRi system was assayed by GFP knockdown as described in (1) with adaptations for *A. baumannii*. Briefly, initial cultures (n=4) were grown from single colonies to saturation (18 h) in 300 µl LB + gent in a deep 96-well plate. These cultures were serially diluted 1:10,000 into 300 µl LB medium with no antibiotic and 0 to 1 mM IPTG and grown back to saturation (15 h). Pelleted cells were resuspended in 300µl 1X PBS and 150 µl was transferred to a clear-bottom black microtiter plate and cell density was determined by OD600, and fluorescence was measured by excitation/emission at 482/515 nm using a Tecan Infinite 200 Pro M Plex plate reader. Fluorescence values were normalized to cell density and to measurements from strains not expressing GFP. This assay was repeated 3 times with 4 independent colonies.

**Stability assay.** Six cultures of *A. baumannii* 19606 *att*Tn*7*::Mobile-CRISPRi (sJMP6335) were serially diluted 1:10^4^ in LB (no selection) and grown to saturation. These cultures were passaged every 24 h for 8 days with each passage being ~13 doublings for a total of ~104 doublings. The final culture was serially 1:10 diluted and 3 µl was spotted on plates with and without gent selection. Also, 50µl of culture was streaked on LB to obtain 44 isolated colonies that were then patched on plates with and without gent selection. No difference in plating efficiency or gent resistance was seen indicating that the Mobile-CRISPRi system remains stably incorporated on the *A. baumannii* chromosome for >100 generations even in the absence of selection.

**Tn*7* insertion location.** Insertion of the CRISPRi expression cassette into the Tn7 att site downstream from *glmS* in *A. baumannii* was confirmed by PCR with primers oJMP60 and oJMP398 (within CRISPRi transposon and upstream of insertion site), oJMP566 and oJMP399 (downstream from insertion site and within the CRISPRi transposon), and oJMP398 and oJMP399 (flanking the insertion site). See Fig S1A.

**GO593_00515 knockdown lysis and plaque assay.** *A. baumannii* strains with chromosomally located CRISPRi expression cassettes and *E. coli* MG1655 (sJMP163) were grown to saturation in liquid culture medium with or without antibiotic selection, respectively. Cultures were diluted 1:200 into nonselective medium and grown ~4 generations to mid-log. *A. baumannii* cultures were then diluted 1:10 into medium with 1 mM IPTG. P1 lysate was added to sJMP163 cultures as previously described as a lysis positive control (5). *A. baumannii* lysates were collected at 7 hours after induction and filter sterilized. Lysates were spotted on bacterial lawns of *A. baumannii* strain ATCC17978 (sJMP4002) to test for phage activity as previously described (5).

**Prophage deletion construction and growth.** We hypothesized that gene G0593_00515 is essential because it is repressing expression of toxic gene(s) in the predicted prophage (GenBank CP046654.1; 84,589-131,937). We should, therefore, be able to select for deletion of the prophage in an *A. baumannii* strain with a CRISPRi knockdown of G0593_00515 because that would receive the toxicity. We used a two-step homologous recombination selection/counterselection approach to delete the prophage. First, we assembled homologous regions (~1 kb) upstream and downstream of the prophage (amplified by PCR with oJMP1568/1569 and oJMP1570/1571) adjacently with the AscI-EcoRI fragment of the R6K ori (pir-dependent) plasmid pJMP1183 which also contains kanamycin resistance and mRFP expression cassettes. This plasmid (pJMP4345) was transferred to the chromosome of an *A. baumannii* GO593_00515 CRISPRi knockdown strain (sJMP4341) by conjugation followed by homologous recombination with selection on LB + kan. Red colonies indicate that single-crossover mutants with the plasmid recombined into the genome were selected and grown to saturation in liquid medium. Second, we counter-selected against presence of the prophage by plating on LB with 1 mM IPTG to induce the CRISPR system. The GO593_00515 knockdown has little detectable growth in inducer, so white colonies retaining the ability to grow should be prophage deletions where a second recombination event removed the integrated vector as well as the prophage. Eight individual isolates (strains sJMP4358-4366) were confirmed by PCR with oligos oJMP1241 and oJMP1242 (within GO593_00515) and oJMP1381 and oJMP1382 (flanking prophage).

### **Prophage deletion/GO593_00515 growth curves.** *A. baumannii* strains with chromosomally located CRISPRi expression cassettes with a non-targeting or GO593_00515-targeting guide with and without deleted prophage (sJMP4324, sJMP4341, sJMP4358 and sJMP4360-4365, respectively) were grown to saturation in liquid culture medium with antibiotic selection. The cultures were diluted 1:200 (OD_600_ ~0.02) into nonselective medium with 0 or 1 mM IPTG and grown for 18 hours in a plate reader (Tecan Infinite 200 Pro M Plex) at 37°C with shaking; growth was measured as OD_600_.

**Analysis of growth phenotypes in strains expressing individual sgRNAs.** Growth of *A. baumannii* strains with chromosomally located CRISPRi expression cassettes (3-6 individual isolates) was analyzed in a plate reader (Tecan Infinite 200 Pro M Plex at 37°C with orbital shaking) in the presence of IPTG inducer and antibiotics. For strains with sgRNAs targeting *nuoH*, *nuoB*, *lpxC*, and a non-targeting control (sJMP10073, sJMP10074, sJMP10076, and sJMP10081, respectively), after growth to saturation in liquid medium with antibiotic selection, cultures were diluted 1:1000 in media with 0.5 mM IPTG and grown for 18 hours prior to dilution 1:1000 into media with 0.5 mM IPTG plus the following antibiotics (in µg/ml): colistin (2 or 4) or rifampicin (0.48 or 0.96), or no antibiotic control and monitoring of growth for 18 hrs. For strains with sgRNAs targeting *murA*, *glnS*, and a non-targeting control (sJMP10072, sJMP10079, and sJMP10080, respectively), after growth to saturation from individual colonies in liquid medium with antibiotic selection, cultures were diluted 1:200 in media with 1mM IPTG and grown ~4 generations to mid-log phase prior to dilution 1:10 into media with 1mM IPTG plus the following antibiotics (in µg/ml): imipenem (0.09), meropenem (0.17), or no antibiotic control and monitoring of growth for 18 hrs.

**Checkerboard assays.** Checkerboard assays were performed to assess the synergy between antibiotic pairs. Two-fold serial dilutions of antibiotics were set up in an 8x8 grid in a 96 well assay plate (Corning 3370) at 2X the final concentration in LB in a final volume of 75 µl. For assays with *A. baumannii* 19606 WT (sJMP490), a saturated culture in LB was diluted to OD600 = 0.2 and 75 µl was added to the antibiotic plate (final volume 150 µl, OD600 = 0.1, and antibiotic concentrations as follows (in µg/ml): fosfomycin (39-2500), imipenem (0.0125-8), meropenem (0.025-16), in addition to no antibiotic controls. Plates were covered with aeroseal and incubated at 37°C, 900 rpm, 24h. For assays with *A. baumannii* Mobile-CRISPRi *nuoH* and *lpxC* knockdown and non-targeting strains (sJMP10074, sJMP10076, and sJMP10081, respectively), plates were set up as above except with 200 µl final volume with a final concentration of 1mM IPTG and antibiotics colistin (0.09375-6), and rifampicin (0.0025-2.56), in addition to no antibiotic controls and shaking at 600 rpm. OD600 of 10% or less than the plate maximum is considered to have been inhibited in that well. Fractional inhibitory concentration (FIC) was determined using the formula: A/MICA + B/MICB = FICA + FICB=FIC index.

#### **Plots.** Data were parsed and visualized using the *tidyverse* collection of packages and plotted using *ggplot2* to generate Sankey plots, volcano plots, and bubble plots. Heatmaps were generated using the *pheatmap* *R* package. Custom plotting scripts can be found in the project GitHub repo.

**SI Methods References**
